# Impact of Mycotoxins and Glyphosate residue on the Gut Microbiome and Resistome of European Fallow Deer

**DOI:** 10.1101/2024.12.05.626988

**Authors:** Adrienn Gréta Tóth, Sára Ágnes Nagy, István Lakatos, Norbert Solymosi, Anikó Stágel, Melinda Paholcsek, Katalin Posta, Szilamér Ferenczi, Zsuzsanna Szőke

## Abstract

Mycotoxins and herbicides residue significant threats to animal health. This study investigated the impact of these toxins on the gut microbiome of the fallow deer, a valuable game species. We analyzed the intestinal contents of deer exposed to varying levels of zearalenone (ZEA) and other mycotoxins and toxic compounds such as aflatoxins, deoxynivalenol, fumonisin B1 and glyphosate residues. Metagenomic analysis revealed significant alterations in the bacterial community composition, particularly in the abundance of specific taxa. Higher ZEA levels were associated with decreased alpha diversity, whereas higher aflatoxin levels had the opposite effect. We also observed changes in the abundance of antibiotic resistance genes (ARGs), suggesting a potential link between mycotoxin exposure and antimicrobial resistance. Furthermore, five complete bacterial genomes were assembled from the metagenomic data. These findings highlight the complex interplay between environmental toxins, the gut microbiota, and animal health. Understanding these interactions is crucial for developing strategies to mitigate the negative effects of toxin exposure on wildlife populations.

## Introduction

Mycotoxins are toxic substances produced by certain moulds in the grains that provide our food and feed. Contaminated grains are a concern to the food production sector worldwide^1^. As climate change progresses, the distribution and toxin-producing capacity of mycotoxin-producing fungi may also change compared to currently known conditions^2^. In recent years, aflatoxins have become an emerging concern in Hungary. Climate change is significantly impacting the distribution and toxigenic potential of *Aspergillus* molds, particularly *A. flavus* and *A. parasiticus*, which are the main producers of aflatoxins^3,4^. Increasing global temperatures, altered precipitation patterns, and increased *CO*_2_ levels are expected to affect the growth and toxin production of these fungi^4^. Notably, *Aspergillus* species and aflatoxins, originally associated with tropical and subtropical climates, are now found in temperate zones^2^.

The deleterious effects of secondary fungal metabolites, commonly known as mycotoxins, have been extensively investigated in both humans and animals. These toxic effects can encompass a wide range, from carcinogenesis and DNA damage to the targeting of specific organs, including the kidneys, liver, intestines, or the nervous and reproductive systems.^1^ Furthermore, co-occurrence, interactions or prolonged exposure and accumulation can alter the toxic levels or the biological effects of mycotoxins.^5^ However, it is not only humans and animals that are affected by the presence of mycotoxins. Bacteria that colonise host organisms also react to the toxic metabolites.

As a part of the rivalry for natural resources, certain mycotoxins block bacterial populations, such as penicillin, the first antibacterial agent ever discovered.^6,7^ Aflatoxins have been studied for their potential antibacterial properties and effects on antibiotic resistance genes. Studies have shown that aflatoxin B1, B2, G1, and G2 exhibit antibacterial activity against the phytopathogenic bacterium *Pseudomonas savastanoi pv. phaseolicola*^8^. At the same time, some bacterial genera can take part in the biological detoxification of mycotoxins by preventing their formation or degrading existing mycotoxin contamination.^9^ Certain segments of the digestive tract of ruminants are harboured by such anti-mycotoxic bacteria. Lactic acid bacteria that are the members of the rumen microbiota are able to convert mycotoxins to less toxic or non-toxic compounds.^10,11^ In addition, several compounds present in the rumen-reticulum compartment are able to bind mycotoxins, that make them unavailable for absorption in the gastro-intestinal tract of the host. For this reason, ruminants are considered less susceptible to mycotoxins than monogastrics.^12,13^ However, a number of mycotoxins can resist rumen degradation, and the relative abundance of the colonising bacterial taxa can change.^14^ Furthermore, ruminants, mostly domestic, captive or supplementary fed, with complex, often silage-rich diets, are exposed to a wide variety of mycotoxins, such as aflatoxins (AFs), zearalenone (ZEA), fumonisin B1 (FB1) or deoxynivalenol (DON) over long periods of exposure.^15^ For the above reasons, the ingestion of mycotoxins has a major impact on the health of ruminants and the economic implications are not negligible.^13^ Additionally, ruminants are susceptible to exposure to further potentially toxic compounds, including those used as herbicides in intensive agricultural settings during feed production. Glyphosate is a nonselective systemic herbicide that inhibits 5-enolpyruvylshikimate-3-phosphate (EPSP) synthase of plants and some microorganisms, including some bacteria but not human or other animal cells. The intake of glyphosate into the rumen of the animals might impair the EPSP pathway of the commensal microbiota.^16^ While the effects of mycotoxin and toxin exposure in domestic ruminants, particularly dairy cattle^17^, have been extensively studied, much less information is available for game species. The investigation of wild animal microbiomes is of critical importance, as these species function as bioindicators, facilitating the comprehension of environmental change impacts and identification of potential human health risks therefore the toxin-associated gut bacteriome study of the fallow deer (*Dama dama*) is presented.

The objective of the study was to perform the shotgun sequencing-based metagenomic analysis of the fallow deer intestinal bacteriome, focusing on toxin-associated changes in the bacterial abundance, diversity and genomics. To assess the genomic effects of the toxins, the set of antimicrobial resistance genes (ARGs), the resistome, was also aimed to be investigated.

## Methods

### Sample collection

The hunts were conducted during the regular hunting season of December–January 2020/2021. Sampling was carried out near Tamási, Hungary, at the forested areas of the South Transdanubian Hills. The principal climatic conditions, the environmental factors and the main game management characteristics of the sampling areas were similar. The sampling areas were all situated within the continental climate zone, characterised by an average precipitation of 500–650 mm with unequal dispersion and an average annual temperature of 9.5 to 11.5 °C. The areas are covered with oak (*Quercus robur* and *Q. cerris*) and black locust (*Robinia pseudoacacia*) dominated, broad-leaved forests surrounded by agricultural lands with maize, wheat, sunflower, and alfalfa. Fallow deers were the dominant game species in all sampling areas, with trophy hunting representing the principal game management goal. To support good quality populations for trophy hunting and vension production, the second most significant fallow deer management objective, intensive methods with strict population control, habitat and game field management and supplementary feeding from autumn to spring are implemented.

After being hunted, the subjects were sampled for this microbiome analysis. Based on the zearalenone (ZEA) toxin levels measured in the gut contents, three study groups of five individuals each were formed; one with low ZEA levels (<10 ng/g), one with medium ZEA levels (10-30 ng/g) and one with high ZEA levels (>30 ng/g). The weight of each sampled individual was measured. The pathological dissection of each individual was conducted and each torso (body without the head, neck, limbs and tail) was weighted.

### Toxin analysis

Mycotoxin measurements were performed on the intestinal content of the hunted fallow deer. Mycotoxin levels were measured using ELISA as was described previously.^18–20^ ZEA, FB1, DON, and total AFs (B1, B2, G1, G2) were assessed through immunoassays. The ABRAXIS® Glyphosate Plate ELISA Kit (PN 500205) was used (Gold Standard Diagnostics, Warminster, US). This kit provides a reliable and accurate method for detecting glyphosate, a commonly used herbicide, in serum samples. To prepare serum samples for analysis, 500 µL of each sample was filtered using a Millipore Amicon centrifugal filter unit. The samples were then centrifuged at 8,000 x g for 15 min to separate the supernatant from any solid particles or debris. After centrifugation, 300 µL of the supernatant was transferred to a microcentrifuge tube that was labeled appropriately. To extract glyphosate from the sample, 200 µL ethyl acetate was added to the tube. The tube was then vortexed for 30 s to ensure thorough mixing of components. Following vortexing, the tube was centrifuged for 3 minutes at 8,000 x g to separate the different phases in the sample. The bottom aqueous phase, which contained glyphosate, was carefully transferred to a new microcentrifuge tube labeled appropriately. The extracted aqueous phase is now ready to be analyzed as a sample using the ABRAXIS® Glyphosate Plate ELISA Kit. Specific instructions for derivatizing the standards, controls, and samples can be found in the Test Preparation section of the kit’s user guide.^21^

### DNA extraction, library preparation and sequencing

DNA was extracted from five intestinal content samples from each three category (low, medium, high ZEA levels) without pooling. DNA extraction was performed using the DNeasy® PowerSoil® Pro Kit (Qiagen, Germany) following the manufacturer’s instructions. Minor modifications were made to optimize the DNA extraction. In brief, 750 l of fecal sample supernatant was centrifuged at 21,000 × g for 5 minutes. The pellet was dissolved in Solution CD1, transferred into PowerBead Pro Tubes (Qiagen, Germany) and incubated at 65°C for 10 minutes. With the use of a MagNA Lyser Instrument (Roche Applied Sciences, Germany), samples were lysed twice at 6,000 × RPM for 30 seconds each. Finally, 70 l of Solution C6 was added and incubated at room temperature for 5 min before centrifugation. DNA concentrations were determined fluorometrically using a Qubit® Fluorometric Quantitation HS dsDNA Assay kit on a Qubit® 4.0 Fluorometer (Thermo Fisher Scientific, USA). The DNA was fragmented and 5’ Adapter:5’-AGATCGGAAGAGCGTCGTGTAGGGAAAGAGTGTAGATCTCGGTGGTCGCCGTATCATT-3’, and 3’ Adapter: 5’-GATCGGAAGAGCACACGTCTGAACTCCAGTCACGGATGACTATCTCGTATGCCGTCTTCTGCTTG-3’ adapters were used. Shotgun sequencing was conducted on an Illumina NovaSeq 6000 instrument (Illumina, USA) with a 150-bp paired-end sequencing run at Novogene Co. Ltd, China. The sequencing yielded a minimum of 20 million reads per sample. To ensure the availability of 20 million reads per sample for shotgun sequencing, we achieved this by re-isolating from each sample until obtaining the required purity OD260/280=1.8-2.0 and concentration 10 ng/L for each metagenomic isolate.

### Bioinformatic analysis

The bioinformatic analyis steps were carried out in two phases. First, for the taxonomic analysis of the paired-end reads, forward and reverse reads were merged with PEAR (v0.9.11).^22^ Quality-based filtering and trimming of the merged reads was performed by TrimGalore (v.0.6.7, https://github.com/FelixKrueger/TrimGalore, accessed on 02/05/2024), setting 20 as a quality threshold and a minimal length of 50 bp. The trimmed reads were dereplicated with VSEARCH (v2.18.0).^23^ The remaining reads were taxonomically classified using Kraken2 (v2.1.3)^24^ using the nt Database (accessed on 03/01/2024) inclusive of GenBank^25^, RefSeq^26^, TPA^27^ and PDB^28^. For this taxon assignment, the–confidence 0.5 parameter was used to obtain more precise species-level hits. The taxon classification data were managed in R (v4.1.2)^29^ using functions of the packages phyloseq (v1.38.0)^30^ and microbiome (v1.16.0)^31^. Prior to contig assembly, only trimming and filtering of the paired-end reads was performed using the same methods and settings as described above. These trimmed and filtered reads were assembled to contigs by MEGAHIT (v1.2.9)^32^ using default settings. The contigs were also classified taxonomically by Kraken2 with the same database as above. All possible open reading frames (ORFs) were gathered by Prodigal (v2.6.3)^33^ from the contigs. The protein-translated ORFs were aligned to the sequences of the Comprehensive Antibiotic Resistance Database (CARD, v.3.2.9)^34,35^ by Resistance Gene Identifier (RGI, v6.0.3) with Diamond^36^. The ORFs classified as perfect or strict were further filtered with 90% identity and 90% coverage. Based on the methods of Hendriksen et al.^37^, ARG abundance was expressed as fragments per kilobase per million fragments (FPKM) of contigs containing ARGs. For the *i*th contig *FPKM_i_* = *q_i_/*(*l_i_* × *Q*) × 10^6^, where *q_i_* is the number of reads that mapped to the contig, *l_i_* is the length of contig and *Q* is the total number of mapped reads. To calculate *q* values, all trimmed and filtered reads were aligned to the contigs by Bowtie (v2.5.3) with the parameter of –very-sensitive-local^38^. All data management procedures, analyses and plottings were performed in R environment (v4.1.2)^29^.

To reconstruct complete bacterial genomes from the metagenomic data set, the binning of the assembled contigs was performed using the following binning tools: SemiBin2 (v2.0.2)^39^, MetaBAT2 (v2.12.1)^40^, MaxBin2 (v2.2.7)^41^ and MetaDecoder(v1.0.18)^42^. The bins were optimised by DAS Tool (v1.1.6)^43^. The taxonomic classifications of the resulting bins were carried out using GTDB-Tk (v2.3.2)^44^ with the Genome Taxonomy Database (GTDB)^45^ and Kraken2 (v2.1.3)^24^, which was run on a database of complete bacterial genomes from National Center for Biotechnology Information (NCBI) RefSeq^26^ built on 05/12/2023. The quality of the genome bins was assessed using CheckM2 (v1.0.1)^46^ and bins with less than 90% completeness were filtered out. For further characterisation of the binned, pre-classified contigs, PGAP (v2024-04-27.build7426)^47^ was used in taxcheck-only mode.

### Statistical analysis

The ARG abundance measures (FPKM of contigs containing ARGs and ARG numbers) were compared between the conditions (mycotoxins, other chemical compounds, weight) using linear models. The within-subject (*α*) diversity was assessed using the numbers of observed genera (richness) and the Inverse Simpson’s Index (evenness). These indices were calculated in 1,000 iterations of rarefied operational taxonomic unit (OTU) tables with a sequencing depth of 1448710. The average over the iterations was taken for each sample. The *α*-diversity expressed by Inverse Simpson’s Index was compared between the ZEA level groups and between the conditions using linear models. The between-subject diversity (*β* -diversity) was assessed by Bray-Curtis distance^48^ based on the relative abundances of bacterial species. Using this measure, principal coordinate analysis (PCoA) ordination was applied to visualize the samples’ variance. The abundance differences in core bacteriome between groups were analyzed by a negative binomial generalized model of DESeq2 package^49^ in R^29^. This approach was applied following the recommendation of Weiss et al.^50^. According to the multiple comparisons, the FDR-adjusted P-value less than 0.05 was considered significant. The statistical tests were two-sided.

## Results

In the results of our study, we first present the most important results of dissection and the toxin analysis, followed by the associations between ARG measures (FPKM, count) and the conditions studied. Following the within- and between-subject diversity of the total bacteriome, the differentiating species of the core bacteriome are presented in relation to the conditions. Since the groups of fallow deers were formed based on the ZEA levels, the statystical tests were performed to compare the ZEA groups and to present the concentration-associated results of all other conditions tested. The set of ARGs with the affected drug classes and the mechanisms of resistance can be seen on Supplementary figure 5 and 6, respectively.

### Dissection and toxin analysis

The results of the toxin analysis and a summary of the most significant pathological findings, if any, derived from the dissections are presented in Table 1 and 2.

**Table 1.** The most important characteristics and pathological findings of the sampled fallow deer. The torso weight includes the body without the head, neck, limbs and tail. CG: Current Gestation, GP: Gestatition Previously, NG: No Gestation.

| ZEA level | Deer ID | Torso (kg) | Age | Sex | Gestation status | Pathology |
| --- | --- | --- | --- | --- | --- | --- |
| Low | TA/4 | 38.7 | Young | Female | CG | - |
| Low | TA/6 | 29.7 | Middle-aged | Female | CG | - |
| Low | TA/9 | 30.5 | Young | Female | CG | - |
| Low | TA/11 | 32 | Middle-aged | Female | CG | - |
| Low | TA/12 | 25.1 | Young | Female | CG | - |
| Medium | TA/1 | 32.2 | Middle-aged | Female | CG | - |
| Medium | TA/3 | 29.6 | Middle-aged | Female | GP | - |
| Medium | TA/7 | 27.3 | Young | Female | CG | Spotted kidneys |
| Medium | TA/8 | 27.1 | Young | Female | CG | - |
| Medium | TA/21 | 27.4 | Young | Female | NG | Vaginal mass |
| High | TA/13 | 26.7 | Middle-aged | Female | CG | - |
| High | TA/14 | 26.1 | Middle-aged | Female | CG | Hardened liver |
| High | TA/16 | 22.9 | Middle-aged | Female | CG | Spotted kidneys |
| High | TA/17 | 28.4 | Young | Female | CG | - |
| High | TA/22 | 27.3 | Young | Female | CG | Hardened liver |

**Table 2.** Intestinal content toxin concentrations in the sampled fallow deer. Toxin concentrations are shown in ng/g. ND: Not Detectable, NM:Not Measured.

| ZEA level | Deer ID | ZEA | AFs | DON | FB1 | Glypho |
| --- | --- | --- | --- | --- | --- | --- |
| Low | TA/11 | 6.077 | 7.272 | 509.160 | 106.932 | 0.259 |
| Low | TA/12 | 0.338 | 10.224 | 373.200 | NDND | 0.334 |
| Low | TA/4 | 4.371 | 10.290 | 899.280 | 20.800 | 0.207 |
| Low | TA/6 | 2.664 | 7.764 | >0.5 | 153.350 | 0.382 |
| Low | TA/9 | 3.312 | 6.000 | >0.5 | ND | 0.471 |
| Medium | TA/1 | 16.477 | 5.100 | 167.160 | 19.000 | 0.401 |
| Medium | TA/3 | 29.438 | 12.036 | 153.000 | 70.900 | 0.320 |
| Medium | TA/7 | 16.679 | 7.602 | 277.800 | ND | 0.389 |
| Medium | TA/8 | 19.419 | 4.146 | 368.280 | ND | 0.418 |
| Medium | TA/21 | 15.267 | 1.528 | 1212.240 | 17.049 | 0.246 |
| High | TA/13 | 31.024 | 1.670 | >0.5 | ND | 0.390 |
| High | TA/14 | 54.704 | 2.835 | 1143.240 | 13.050 | 0.363 |
| High | TA/16 | 35.945 | 3.451 | 359.520 | 20.000 | NM |
| High | TA/17 | 72.867 | 2.535 | 985.440 | ND | NM |
| High | TA/22 | 81.458 | 1.290 | 1218.720 | 17.336 | 0.381 |

### Analysis of ARG measure variations

The linear model results of the FPKM of contigs containing ARGs and the examined conditions are presented on Table 16. Two conditions were associated with statistically significant effects, with zearalenone and aflatoxin concentrations having a positive and negative effect on FPKM values respectively.

In contract to the FPKM related results, no significant associations were revealed in case of the ARG numbers and the examined conditions (see Table 4).

**Table 3.** Linear model results of ARG FPKMs and units of the affecting conditions.

| Model | $\beta$ | Std. Error | p-value |
| --- | --- | --- | --- |
| FPKM ∼ ZEA_cc | 0.013 | 0.004 | 0.001** |
| FPKM ∼ AFs_cc | -0.096 | 0.020 | 5.941e-06*** |
| FPKM ∼ DON_cc | -1.122e-05 | 1.391e-04 | 0.936 |
| FPKM ∼ FB1_cc | 0.005 | 0.004 | 0.260 |
| FPKM ∼ Glypho_cc | 0.510 | 1.506 | 0.736 |
| FPKM ∼ Weight | -0.074 | 0.042 | 0.086 |

**Table 4.** Linear model results of ARG numbers and units of the affecting conditions.

| Model | $\beta$ | Std. Error | p-value |
| --- | --- | --- | --- |
| n ∼ ZEA_cc | 0.017 | 0.129 | 0.897 |
| n ∼ AFs_cc | -1.027 | 0.921 | 0.289 |
| n ∼ DON_cc | -0.006 | 0.007 | 0.430 |
| n ∼ FB1_cc | -0.014 | 0.014 | 0.346 |
| n ∼ Glypho_cc | 10.704 | 69.250 | 0.881 |
| n ∼ Weight | -0.699 | 1.369 | 0.620 |

The visualization of the associations of FPKM values or ARG numbers and the examined conditions can be seen on Supplementary figure 7-12 and 13-18, respectively.

### Within-subject diversity analysis

The numbers of observed species and Inverse Simpson’s Index -diversity metrics by zearalenone levels are shown in Fig 1. The Inverse Simpson’s Index outlier by high zearalenone concentrations is sample TA17. Alpha diversity showed no significant difference between groups in either metric. By the comparison of high and medium ZEA levels, the P-value was 0.1358, while by medium and low levels the P-value was 0.7612. By high and low ZEA level groups, the comparative metric was P = 0.0855.

**Figure 1.**
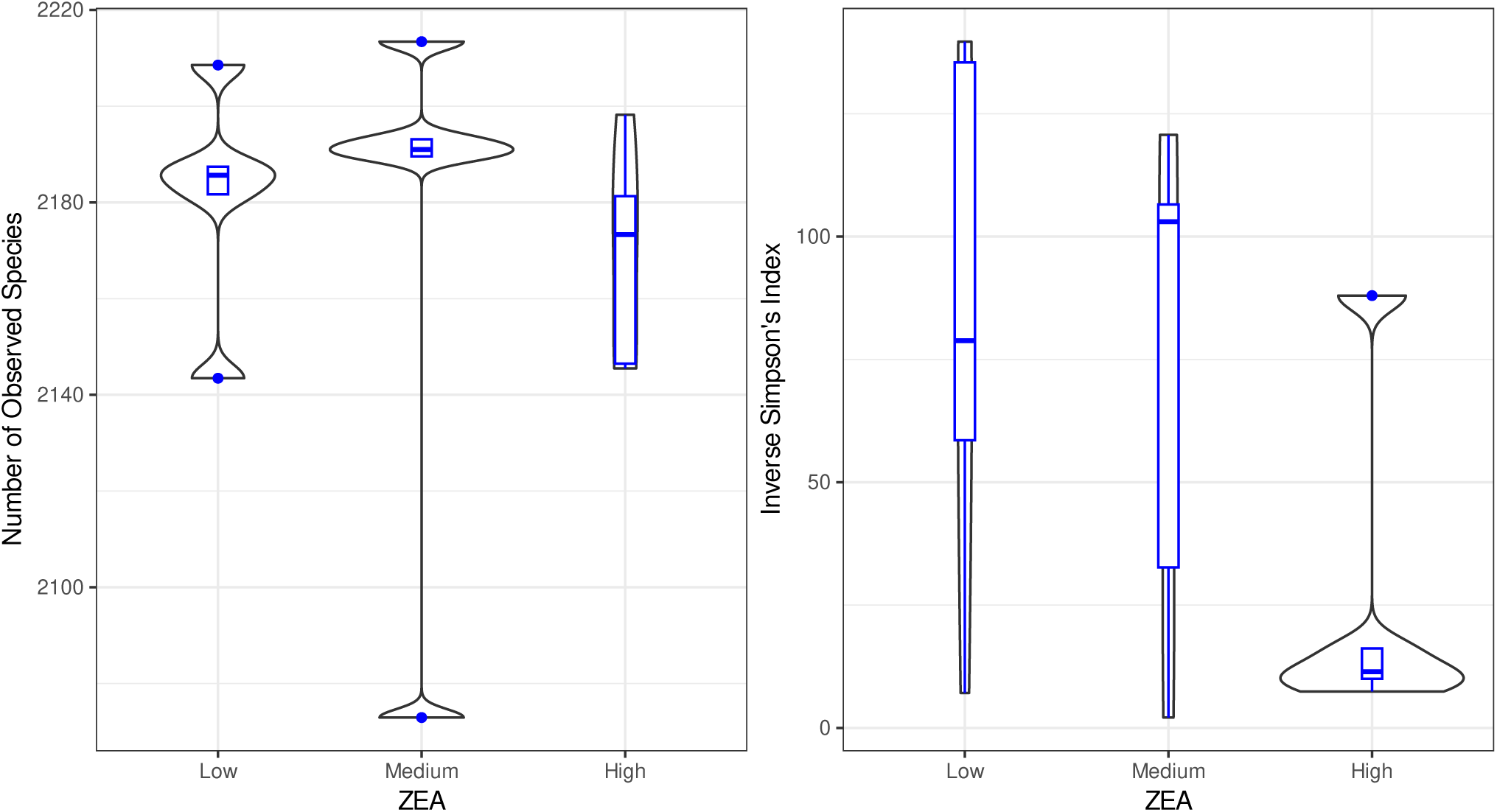
Alpha diversity metrics across different ZEA levels (Low, Medium, High). The left panel displays the number of observed species, while the right panel shows the Inverse Simpson’s Index as a measure of diversity. Violin plots illustrate the distribution of each metric, with embedded box plots indicating median values and interquartile ranges.

The results of *α*-diversity expressed by Inverse Simpson’s Index compared between the conditions using linear models are shown on Table 5. Higher ZEA levels are associated with distinct diversity patterns, with a noticeable reduction in species diversity and evenness at the highest ZEA concentration. In samples with higher AF concentrations, the *α*-diversity was significantly (p = 0.004) higher than in samples with lower aflatoxin concentrations. Furthermore, samples deriving from heavier fallow deers were associated with higher *α*-diversity than samples from lighter animals.

**Table 5.** Linear model results of *α*-diversity expressed by Inverse Simpson’s Index compared between the conditions.

| Model | $\beta$ | Std. Error | p-value |
| --- | --- | --- | --- |
| InvSim ∼ ZEA_cc | -0.697 | 0.526 | 0.208 |
| InvSim ∼ AFs_cc | 10.239 | 2.951 | 0.004** |
| InvSim ∼ DON_cc | -0.043 | 0.029 | 0.163 |
| InvSim ∼ FB1_cc | 0.446 | 0.431 | 0.335 |
| InvSim ∼ Glypho_cc | 3.097 | 215.155 | 0.989 |
| InvSim ∼ Weight | 7.956 | 3.152 | 0.025* |

### Between-subject diversity analysis

The variability of the samples’ genus profiles ( *β* -diversity) is visualized by PCoA ordination (Fig. 2) based on Bray-Curtis distances. The PERMANOVA analysis of the bacterial composition on the genus level revealed no significant difference between the samples originating from the the different ZEA level groups (p = 0.175).

**Figure 2.**
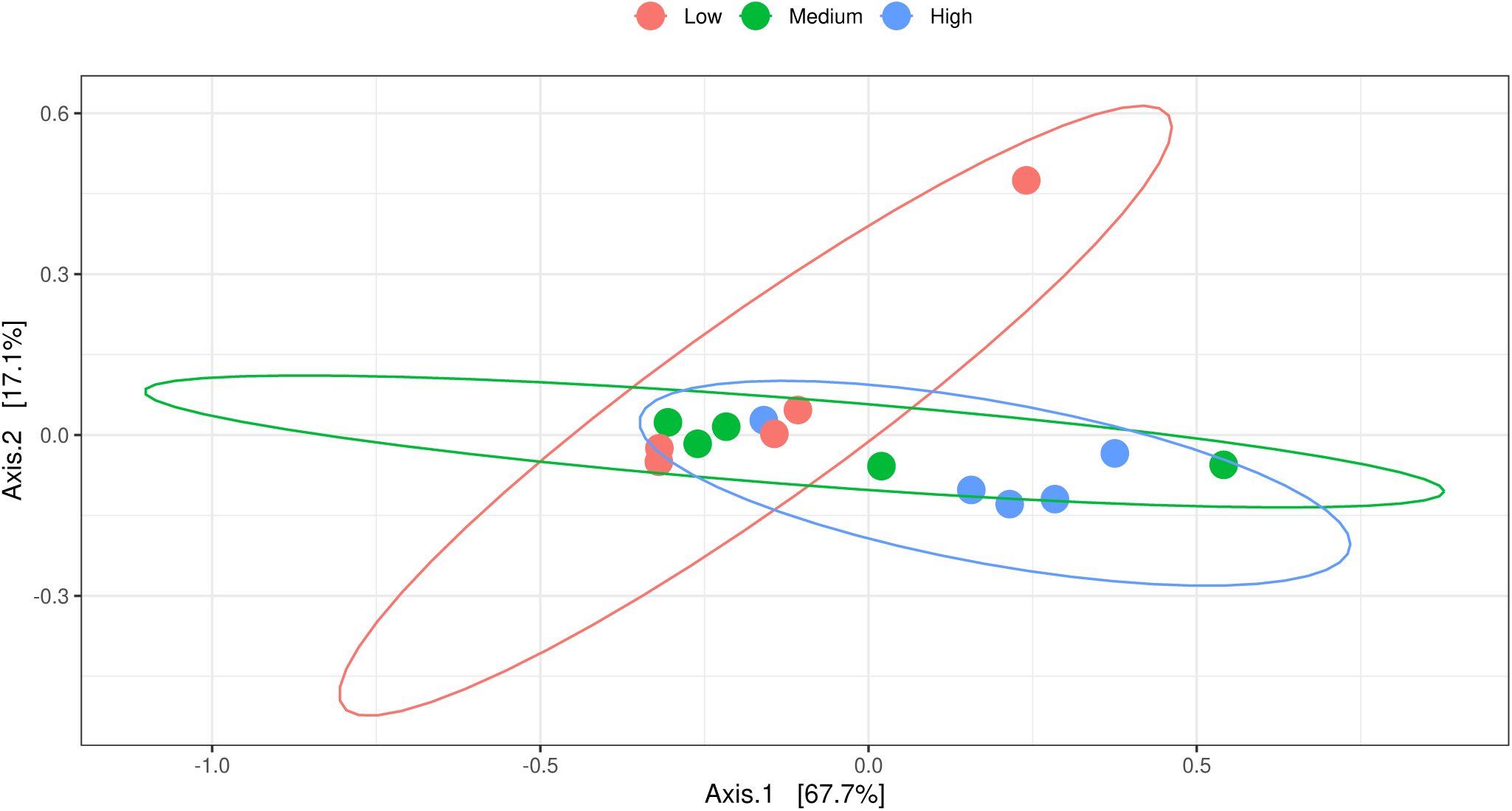
PCoA plots of *β* -diversity estimated based on the bacteriome of fallow deer samples considering the zearalenone levels.

### Core bacteriome analysis

The core bacteriome members having relative genus-level abundance above 0.1% in the samples are presented by zearalenone levels on Fig. 3.

**Figure 3.**
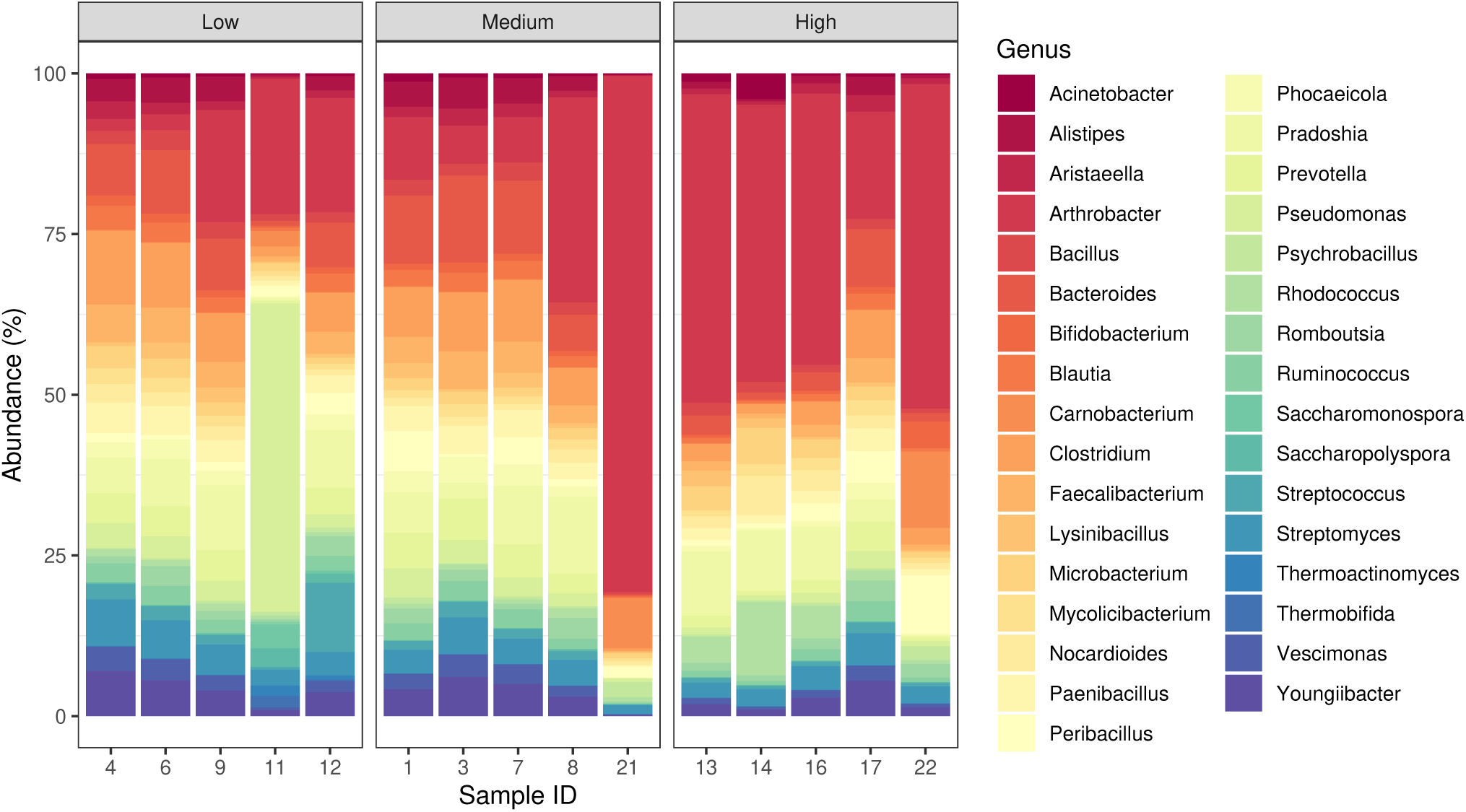
Genus-level core bacteriome composition of the fallow deer samples by zearalenone levels.

The abundance differences (log2 median fold change, Log2FC) of groups per taxon comparison are summarized in Suppl. table 7-9 by zearalenone levels and Suppl. table 10-15 for each examined condition. Comparing low and medium zearalenone levels, the following genera showed significant (adjusted P =< 0.05) differences in abundance: *Saccharopolyspora, Thermoactinomyces*. By the comparison of low and high ZEA levels the genera of *Thermoactinomyces, Rhodococcus, Saccharopolyspora, Acinetobacter* and *Nocardioides* appeared to have significant differences. Between medium and high ZEA levels, *Rhodococcus* and *Nocardioides* abundances were significantly different. The most significant changes in the abundance of the core bacterial genera were found betweenlow and high zearalenone levels (5 taxa). Of these, two-two genera showed significant abundance differences by low to medium and medium to high zearalenone levels. Furthermore, the following genera showed statistically significant differences by the comparison of the examined conditions: by zearalenone concentrations *Rhodococcus, Acinetobacter, Bifidobacterium, Nocardioides* and *Romboutsia*, by aflatoxin concentrations *Saccharomonospora, Rhodococcus, Thermoactinomyces, Arthrobacter, Psychrobacillus, Nocardioides, Blautia, Peribacillus, Carnobacterium, Thermobifida, Streptococcus, Faecalibacterium, Pseudomonas, Acinetobacter, Microbacterium, Saccharopolyspora, Vescimonas, Romboutsia, Alistipes, Lysinibacillus, Phocaeicola* and *Mycolicibacterium*, by deoxinivalenol (DON toxin) *Carnobacterium, Psychrobacillus, Peribacillus, Bifidobacterium, Alistipes, Streptomyces, Mycolicibacterium, Nocardioides, Phocaeicola, Prevotella* and *Bacteroides*, by fumonisin *B*_1_ (FB1) *Saccharomonospora, Thermoactinomyces, Thermobifida, Saccharopolyspora* and *Pseudomonas*, by glyphosate *Saccharomonospora, Thermobifida, Thermoactinomyces, Pseudomonas* and *Saccharopolyspora* and by weight *Pseudomonas, Thermobifida, Arthrobacter, Romboutsia* and *Rhodococcus*. A summary of the above significant ZEA levels, toxin concentrations and weight changes in the core bacteriome are shown in Fig. 4.

**Figure 4.**
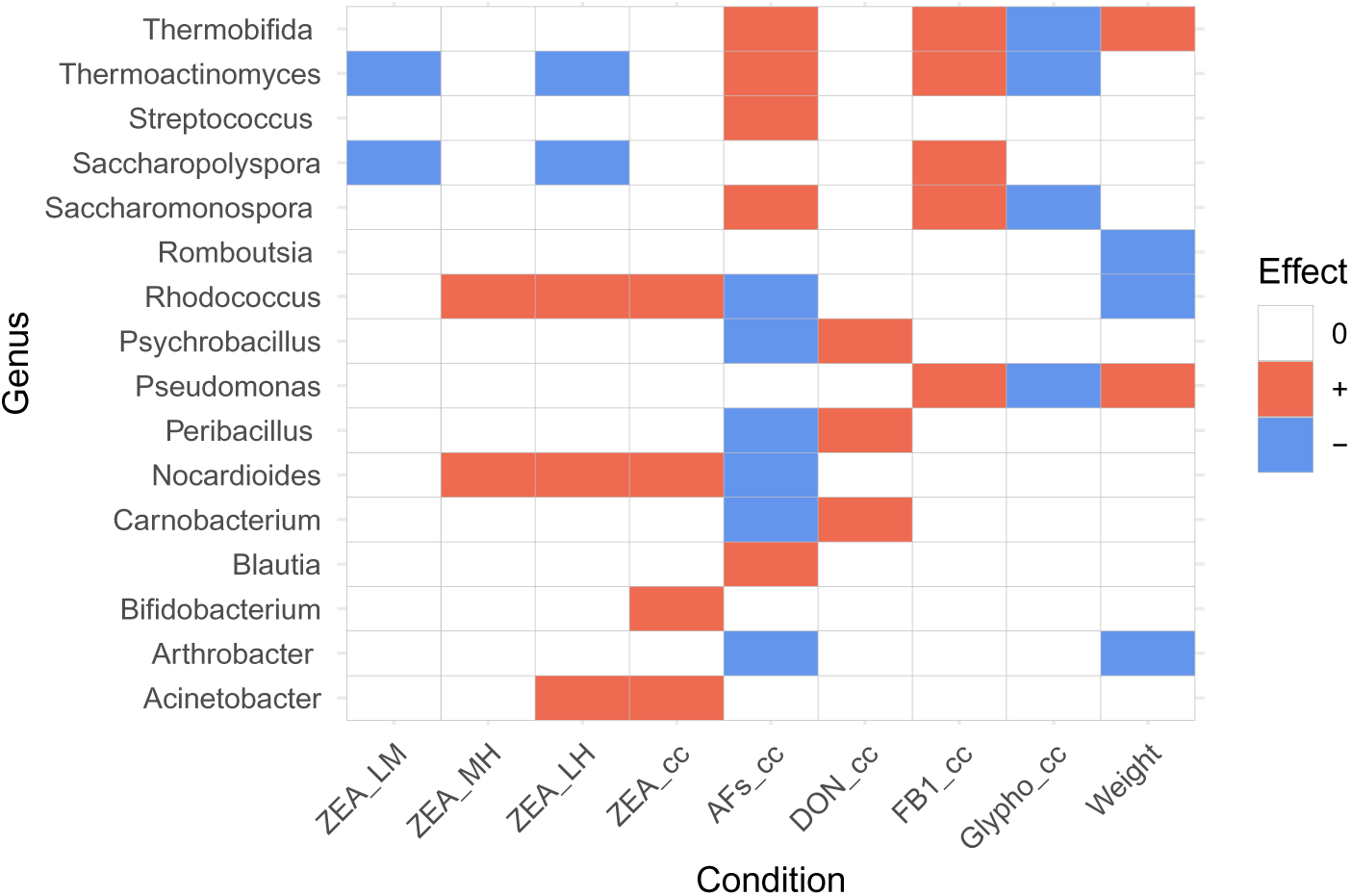
Significant changes in the core bacteriome, considering the tested conditions. In case of ZEA both the ZEA level associated and the ZEA concentration associated results are represented. The colors represent the following: White (0): No significant change, Red (+): Inreased abundance with rise in the toxin level/concentration or weight, Blue (-): Decreased abundance with rise in the toxin level/concentration or weight.

### Complete bacterial genomes

Based on the binning results of the metagenomic data set, 86 potential bacterial genome hits were identified in all samples with a CheckM2^46^ completeness rate of more than 90%. Of these, five bacterial genomes achieved high confidence in the PGAP^47^ analysis. Details of these hits are given in table 6.

**Table 6.** PGAP^47^ characterisation of the bacterial genomes identified with High confidence. ANI : ANI value between this assembly and the type listed in this row. Coverages : query-coverage and subject-coverage of this assembly (query) and the type (subject). NewSeq : the count of bases best assigned to this type assembly. Assembly : NCBI Release-id of the type-assembly.

| Species | Confidence | ANI | Coverages | NewSeq | Assembly |
| --- | --- | --- | --- | --- | --- |
| *Saccharopolyspora rectivirgula* | High | 99.852 | 94.7 97.8 | 2142474 | 1144898 |
| *Streptococcus equinus* ATCC 33317 | High | 98.225 | 90.3 91.1 | 1524505 | 1303318 |
| *Escherichia coli* SE11 | High | 99.289 | 97.3 72.5 | 2462306 | 11188 |
| *Saccharomonospora viridis* | High | 99.745 | 99.0 91.3 | 562898 | 1390408 |
| *Bifidobacterium pseudolongum* subsp. *pseudolongum* | High | 99.235 | 85.4 89.2 | 1432934 | 1189548 |

## Discussion

The health of herbivorous animals can be maintained despite their potentially phytotoxin-and mycotoxin-rich diet through a variety of defensive mechanisms. On the one hand, protection is provided by substances (e.g. enzymes, tannin-binding proteins in saliva) and physiological mechanisms (impermeable intestinal epithelium) of their body cells. On the other hand, the role of symbiotic microorganisms living in the gastrointestinal tract of herbivorous mammals is even more important in neutralising biological agents.^51^ During the evolution of eukaryotic organisms, several solutions have evolved to accommodate the large number of colonizing microorganisms. One of the most obvious examples is the foregut fermentation chambers of ruminants. The microbiota in the gastric compartments are able to detoxify secondary metabolites that enter with the ingested plant material. As a result, the subsequent intestinal sections, where absorption takes place, are under a relatively lower toxin load. This explains the phenomenon that ruminants are more resistant to mycotoxins and phytotoxins.^51^ However, the sensitivity of ruminants to toxins is still considerable due to the high intakes and long, continuous periods of exposure.^13^

At the same time, exposure to mycotoxins or other toxins affects not only the well-being of the host, but also its colonising microbes. The effects on bacteria can be very pronounced. If the environmental conditions alter, bacteria are capable of the uptake of genes from their environment or from eachother to aid their survival.^52^ Thus, toxin-induced changes can appear in the genomic level. Besides certain mycotoxins, such as penicillin, that act as actual antimicrobial agents, the levels of other toxins can also underlie changes in the resistome.^53^ AFs demonstrate antibacterial properties and they may also contribute to antibiotic resistance^54^.

Concurrently, our study revealed that the abundances of ARGs only exhibited statistically significant alterations in conjunction with the changes in the concentrations of ZEA and AF. Given that mycotoxins and certain antibiotics are both bioactive substances produced by fungi, the shared origin can be manifested by the similarity of their chemical structure.^55^. To illustrate, one of the AFs, aflatoxin B1, belongs to the structural class, which includes tetracyclines as well. Zearalenone, meanwhile, has a structure that is similar to some macrolides.^55^. Consequently, it can be inferred that the higher abundance of ARGs observed in samples with elevated ZEA concentrations represents a bacterial response to the presence of the mycotoxin. However, the results were found to be contradictory for high AF concentrations, with the higher concentrations observed to decrease the abundance of the ARGs. The question emerged whether this trend remains when testing changes in different mycotoxin levels and antibiotic resistance genes with chemically similar structures. When the abundance of ARGs against macrolides and tetracyclines was tested by different ZEA and AF concentrations, a significant correlation was found only for the latter pair (see supplementary figures 19-20 and supplementary table 16). Despite the chemical similarity between macrolides and zearalenone, as well as between tetracyclines and aflatoxin, the abundance of the related antibiotic group-specific ARGs follows the trend observed for all ARGs in conjunction with the concentrations of the mycotoxins. It is possible that these findings may be explained by hitherto unexplored fungal-bacterial interactions, which may be synergistic or antagonistic in nature. However, these interactions fall beyond the scope of the present study. Nevertheless, in accordance with the findings of other studies, the levels of mycotoxins had an impact on the ARG pattern of the microbiome^53,56^.

As no significant results were obtained for changes in ARG numbers in response to mycotoxin concentrations, as opposed to ARG abundance, mycotoxins did not appear to be able to induce significant changes in ARG diversity. However, ARG abundance does not necessarily vary with ARG diversity.^57^

Considering the over ARG diversity results, TA13, an approximately 9-year-old doe, that is also the oldest animal involved in the study, exhibited an outstanding number of various ARG types. This finding is consistent with current knowledge regarding the accumulation of ARGs in the human gut microbiota from childhood to adulthood, which demonstrates a progressive increase in complexity with age.^58^

The detoxification capacity of a herbivore is also influenced by the type of herbivory. Browsing animals (as all cervids are in snowy winters^59^) have a diet that allows them to consume more bioactive compounds than grazing ones and are therefore generally more resistant to them.^60^ A possible reason for this may be that a more diverse microbiota confers greater functional resistance to the host organism, promoting more effective coping mechanisms with changes in environmental conditions (e.g. seasons).^61,62^ This may also explain why higher *α*-diversity is associated with higher body weight. A more diverse microbiota is thought to be more efficient at processing a more lignin-rich browsing diet, thereby promoting weight gain in the host. However, de Freitas and colleagues found no association between microbiome diversity and weight gain in beef cattle^63^. The presence and/or abundance of certain taxa may have a greater impact on the productivity of ruminants.^64^.

Furthermore, given the similarities between the detoxification responses to phytotoxins and mycotoxins^51^, it is conceivable that the gut microbiome may respond to high dietary mycotoxin levels in a similar way to the presence of phytotoxins. This could be a possible explanation for the finding that higher *α*-diversity was observed at high AF levels than at lower AF levels. TA17, an Inverse Simpson’s Index outlier with a high ZEA level, was a young animal that, in contrast to the other high ZEA level animals, exhibited no pathological changes in the liver or kidneys. However, the intestinal ZEA level was among the highest in the category with high ZEA levels, and the liver concentration of alpha-zearalenol (alpha-ZOL), a derivative of ZEA was also considerable, as reported by Lakatos and colleagues.^18^

In light of the tests of toxin concentrations and weight relative to the abundance measures of ARGs and bacterial *α*-diversity, no statistically significant changes were observed in any case in response to DON, FB1 and glyphosate. In contrast to this, DON has been shown to exert direct damage to the intestinal barrier function and contribute to the dysbiosis of the intestines in several species.**^?^**^,65–67^ In poulty, indices of *α*-diversity including Chao, ACE and Shannon were all fond to be reduced in groups treated with the toxin.^65^ Similarly, *α*- and *β* -diversity were observed to decrease in mice in the presence of DON.^66^ In another study, DON altered the abundance of microbial communities in dairy cows, thereby detrimentally affecting rumen function and potentially threatening production performance and the overall health of cows. The negative effects were found to be even more pronounced by cows with a high-starch diet.^67^ The relatively low-starch diet of the fallow deer may have contributed to the finding that DON has had negligible effects on the microbiome.

Like DON, FB1 has also been shown to alter the diversity and composition of the fecal microbiota in mice.^68^ In addition to dysbiosis, FB1 has also been described to impair intestinal epithelial barrier function in humans.^69^ Similarly, Hartinger and colleagues reported that FB1 severely impacted the *β* -diversity structure of cattle.^14^ At the same time, ruminants are more tolerant to FB1 than monogastric animals, despite the poor microbial metabolism in the rumen.^70^ Nevertheless, fumonisins are minimally absorbed by ruminants and excreted in the feces in an unmetabolised form.^71,72^ The reason for the lack of significant changes in the fecal microbiota metrics studied would require further investigation. In the case of glyphosate, our findings are consistent with the existing literature, as other authors have also found no evidence that this compound affects microbial diversity or abundance of microbial taxa at any concentration.^73^

In regard to the *β* -diversity of the three ZEA level groups, our findings were consistent with those of Hartinger and colleagues. Their work on the acute effects of ZEA and fumonisin toxins in dairy cows also demonstrated that FB1 has a significant effect on ruminal *β* -diversity, whereas ZEA has no significant effect on it. The effect of FB1 can be explained by its antibacterial activity against certain bacteria, thus reducing their abundance and providing an advantage to other species. Although there was no significant change in *β* -diversity in the presence of ZEA, the abundance of certain genera was altered. This also suggests that mycotoxins can adversely affect the host microbiome, causing health problems.^14^

The similarity of the mycotoxins to antibiotics may also elucidate why the abundance of members of the core bacteriome was significantly disparate in the three ZEA level groups, mostly comparing the low ZEA level groups to the high ones. The abundance of *Saccharopolyspora spp.* was found to be elevated in groups with higher ZEA levels. Concurrently, this genus has been linked to the synthesis of a range of biologically active compounds, including macrolides.^74^ It is therefore unexpected that the species within this genus do not appear to possess defensive mechanisms that would allow them to resist the impact of zearalenone, which shares structural similarities with macrolides. Furthermore, several species of *Saccharopolyspora* and *Thermoactinomyces*, another significantly affected core bacteriome member, are frequently found in mouldy hay and other fodder^75^. Despite this, the ZEA-associated changes in abundance of these genera cannot be explained by the physical introduction of microbes into the mouldy feed, as their abundance decreased along with the increase of ZEA levels. The abundance of *Rhodococcus spp., Acinetobacter spp.* and *Nocardioides spp.* increased in higher ZEA level groups. In line with this finding, the members of the genus *Rhodococcus* are able to degrade a wide range of xenobiotic substances, including zearalenone^76^. Similarly, certain *Acinetobacter* species are characterized by the presence of multiple enzymes that facilitate the biodegradation of zearalenone.^77^ It is well documented that Nocardioides has the capacity to degrade both DON and aflatoxin.^78^ However, it is also reported that it is capable of degrading zearalenone.^79^ It may therefore be hypothesised that the enhanced abundance of these taxa is due to their capacity to degrade toxins. As illustrated in Fig. 3, the core bacteriome of TA21, a sample of medium ZEA level, exhibits notable differences from the other samples. This is evidenced by the dominance of the microbiota by *Arthrobacter spp.*. As described by Lakatos and colleagues, the dissection revealed a range of pathological findings in the individual fallow deer, including vaginitis, metritis and enteritis. Additionally, the liver exhibited high concentrations of aflatoxin, indicating prolonged exposure to this toxin.^18^ Furthermore, changes in ZEA levels and ZEA concentrations in the core bacteriome were consistent. The highest number of significant changes were found in relation to changes in AFs concentration. At the same time, interestingly, AFs were the only mycotoxins, the rise in the concentration of which decreased the abundance of certain bacterial genera. In line with this, the exposure to various mycotoxins is described to affect some bacteria, including opportunistic pathogens beneficially in ruminants^80^ or in other animal species.^6^ Glyphosate was the only toxic compound whose increase always caused a decrease in the significantly affected genera, including *Pseudomonas spp.*, which can be opportunistic pathogens, or *Thermoactinomyces spp.*, which can be responsible for a disease of One Health relevance, farmer’s lung.^81,82^

The five complete genomes that have been assembled from the shotgun sequenced metagenomic dataset have all been previously associated with ruminants in the scientific literature. The members of thermophilic actinomycetes, *Saccharopolyspora rectivirgula*, formerly known as *Micropolyspora faeni*, and *Saccharomonospora viridis* contributes to the deterioration of hay, grain and straw stored at high moisture levels. *S. rectivirgula* has been associated with equine asthma, formerly known as heaves or recurrent airway obstruction in horses.^83^ In addition, both agent are considered a major cause of extrinsic allergic alveolitis, commonly known as farmer’s lung disease, a form of hypersensitivity pneumonitis in humans which may lead to irreversible lung damage. As such these species are of public health significance.^82,84,85^ Because these species are fastidious, their traditional phenotypic identification is challenging.^86^ *Streptococcus equinus* is a member of the *Streptococcus bovis/Streptococcus equinus* Complex that consists of bacteria, frequently colonizing the rumen, colon, crop, and cloaca of animals and humans.^87^ Despite being predominantly commensal, cases of their commensal-to-pathogens transition have been described, raising attention to their emerging opportunistic pathogenic potential.^88,89^ Certain members of the complex are food fermentative organisms.^90^ Due to their equivalence in several genomic features, their identification at the species level requires the combination of phylogenetic and functional analyses.^87,89^ Thus, species-level taxonomic classification based on genotypic methods alone is disputable. Interestingly, on the American Type Culture Collection (ATCC) Genome Portal (https://genomes.atcc.org, accessed: 14/10/2024)^91^, which is the only authenticated reference genome database for ATCC microbes, the largest general culture collection in the world, the identified strain is referred to as *Streptococcus bovis* Orla-Jensen. Despite being commonly found in the intestines^92^, *Escherichia coli* is reported as a potential health threat in cervids.^93,94^ The closest strain hit, SE11 (O152:H28), however has been described in the gut of healthy adults.^95^ *Bifidobacterium spp.* are one of the most important groups of bacteria in human and animal feces. Moreover, due to their strict nutritional requirements and poor growth outside the gut, they can be useful indicators of fecal pollution.^96^

## Consclusion

Although the results represent a transient state of the toxicological measures of the intestinal content of fallow deer, the microbiome composition and genomic characteristics of these animals have been scarcely studied thus far. In addition to the identification of the taxonomic and genomic properties of the bacteriome, the presence of toxins in the gut content, and thus, presumably, in the diet of the fallow deer, could have been confirmed. Although the ingestion of mycotoxins and other pervasive toxins can be linked to advantageous outcomes, similar to the case of the antibiotic penicillin, more deleterious effects have been observed in association with these compounds. The majority of studies examining the impact of mycotoxins on humans and animals concentrate on the direct effects exerted by these secondary metabolites on the host organisms. However, our focus was on the impact of mycotoxins on the colonising bacteriome of fallow deer. With regard to the health status of the host, the impact on the microbiome is more indirect. Nevertheless, the state of the microbiome is inextricably linked to the well-being of ruminants.^97^ It is therefore of the utmost importance to gain a deep understanding of the genomic aspects of bacterial exposure to mycotoxins and other toxins. Furthermore, the significance of this field is reinforced when considering the potential changes in the number of ARGs. In addition to the potential impact on the efficacy of antibiotics administered to animals, public health concerns may also arise. The presence of mycotoxins may lead to an expansion of the ARG set, increasing the likelihood of horizontal ARG transfer between bacteria. This raises One Health and public health concerns, as the ARGs may be transmitted between the animals and hunters or consumers who are in close physical contact with them.

## Declarations

### Ethics approval and consent to participate

According to the statement of the Institutional Review Board (NAIK MBK MÁB 004-09/2018), the study is not considered as an experiment with animals because the researchers collected samples from legally harvested fallow deer hinds; consequently, the ethical treatment rules are not applicable.

### Consent for publication

Not applicable.

### Data Availability

The raw short-read data are publicly available and accessible through the PRJNA1091038 from the NCBI Sequence Read Archive (SRA).

### Author contributions statement

AGT, NS, SF and ZS take responsibility for the data’s integrity and the data analysis’s accuracy. AGT, NS, SF and ZS conceived the concept of the study. IL collected the biological samples. AS, IL, MP and ZS performed the laboratory processes. AGT and NS participated in the bioinformatic analysis. AGT, NS, SÁN, SF and ZS participated in the drafting of the manuscript. AGT, AS, IL, KP, MP, NS, SÁN, SF and ZS critically revised the manuscript for important intellectual content. All authors read and approved the final manuscript.

## Authors’ information

Not provided.

## Supplementary materials

**Figure 5.**
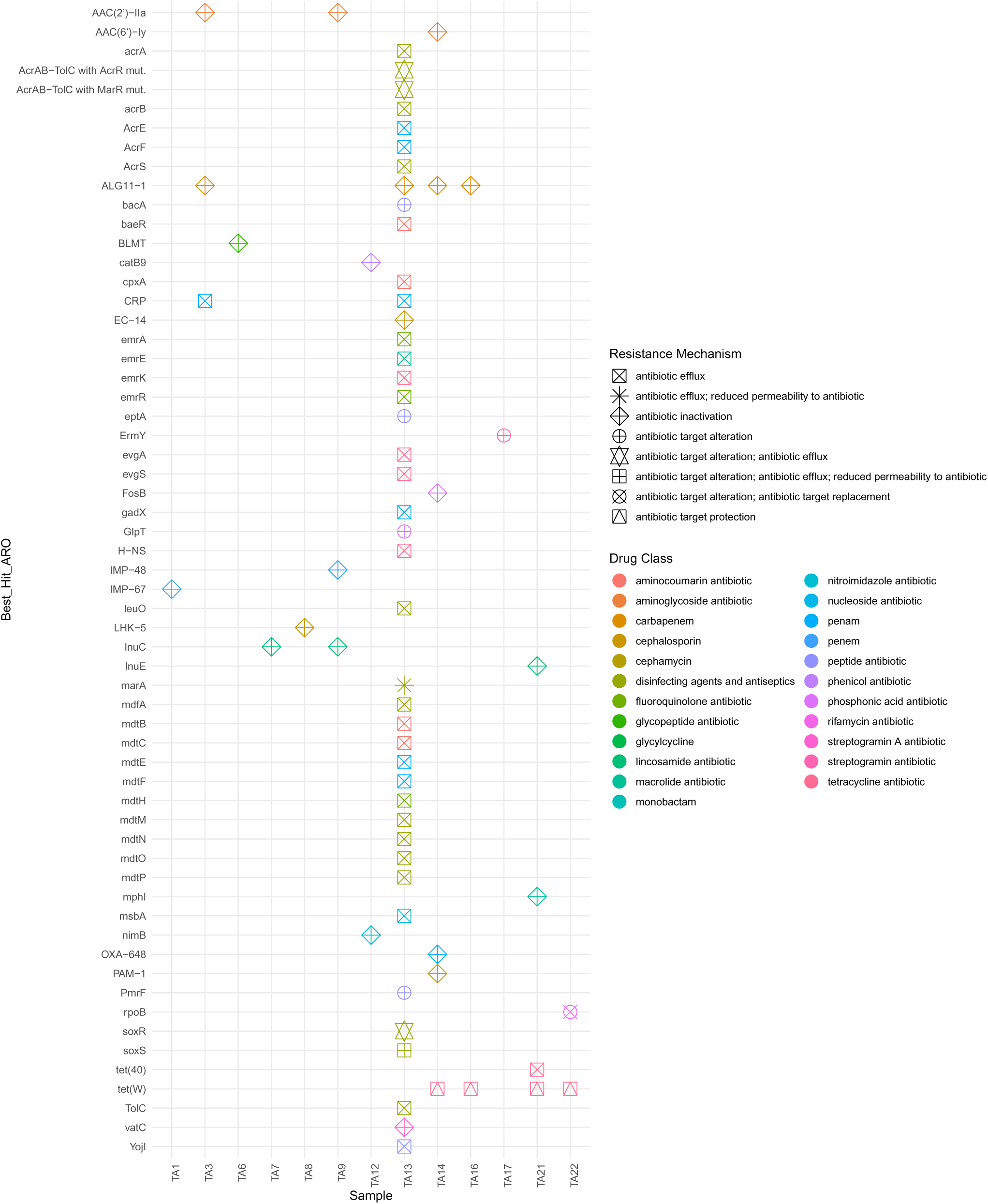
ARGs (Best_Hit_ARO) found in the studied samples. The colors represent the potentially affected drug classes and the shapes indicate the mechanism of AMR.

**Figure 6.**
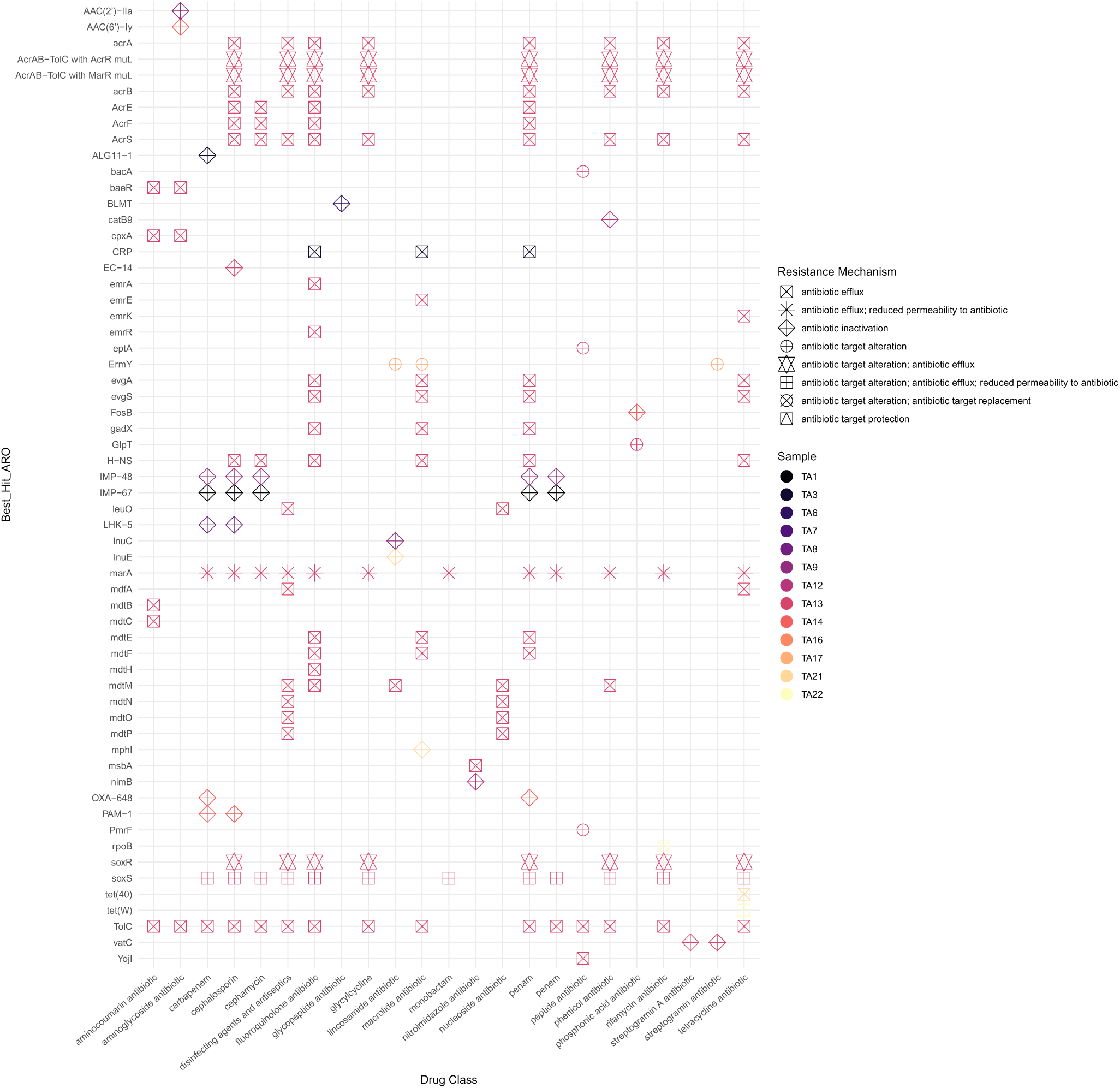
ARGs (Best_Hit_ARO) with the affected drug classes. The colors represent the ARGs’ sample of origin and the shapes indicate the mechanism of AMR.

**Figure 7.**
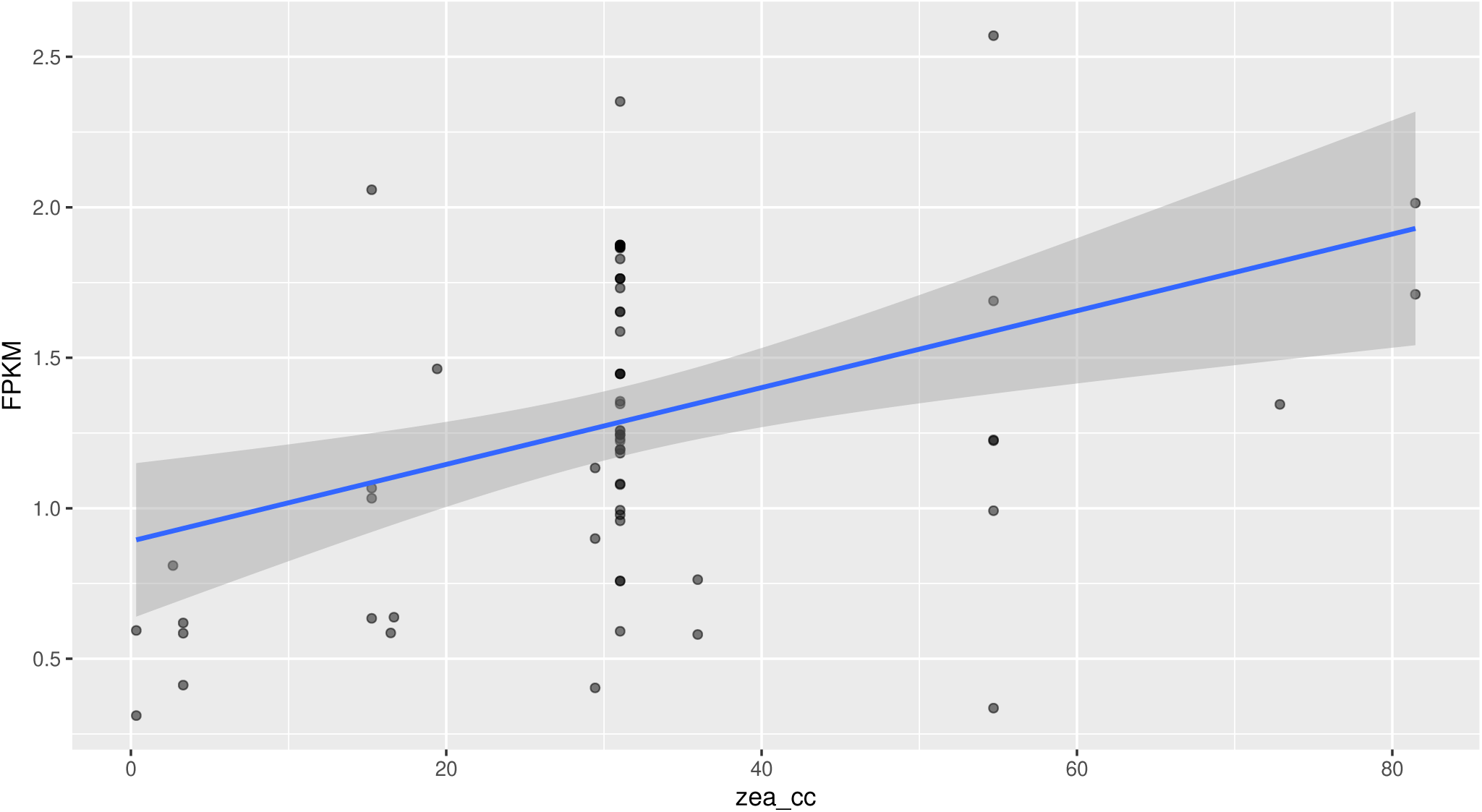
Relationship between ZEA concentration (zea_cc) and ARG counts (FPKM). The scatter plot shows individual data points for ARG FPKM values across a range of ZEA concentrations, with a fitted linear regression line (blue).

**Figure 8.**
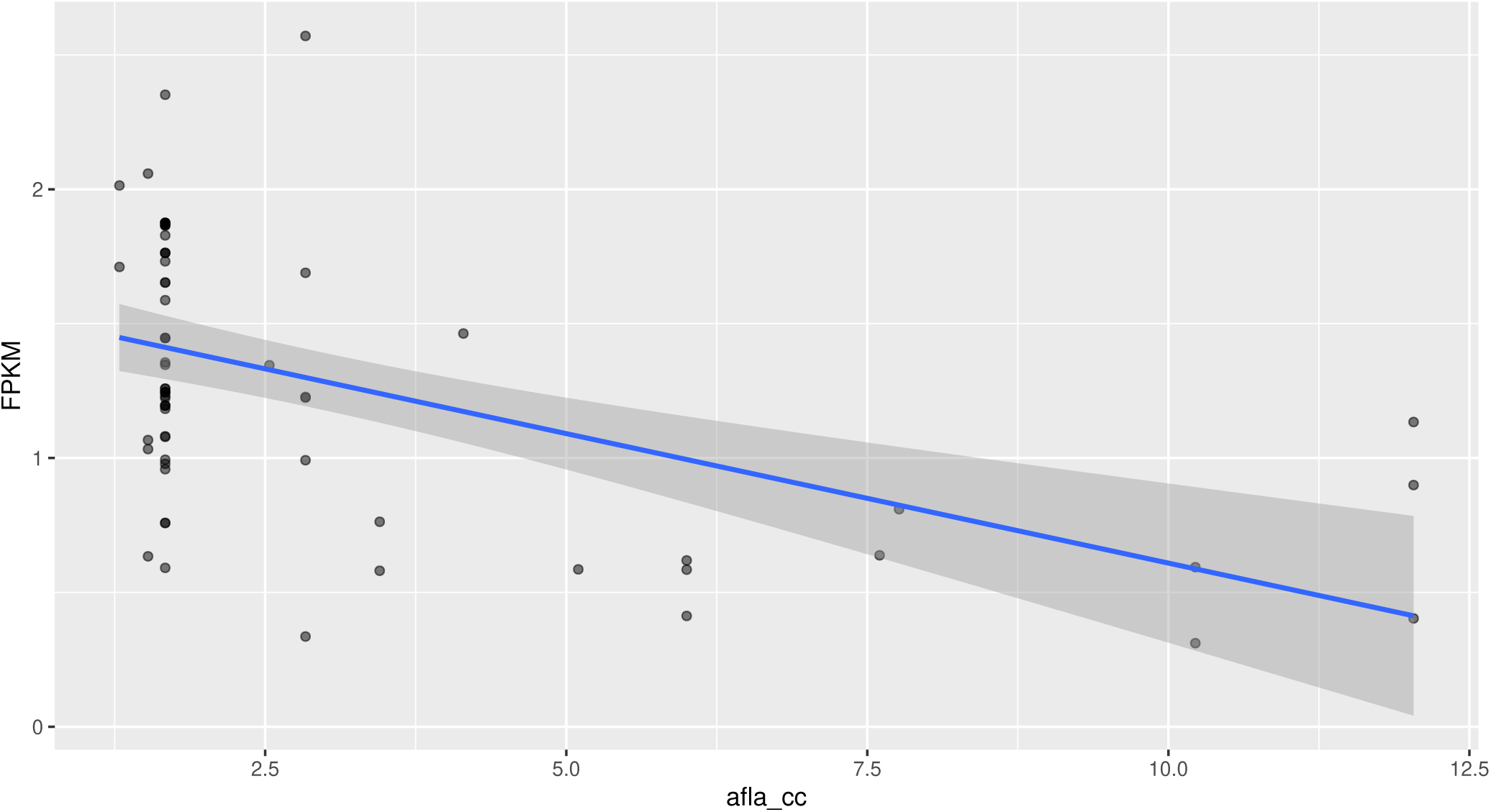
Relationship between AFs concentration (afla_cc) and ARG counts (FPKM). The scatter plot shows individual data points for ARG FPKM values across a range of AFs concentrations, with a fitted linear regression line (blue).

**Figure 9.**
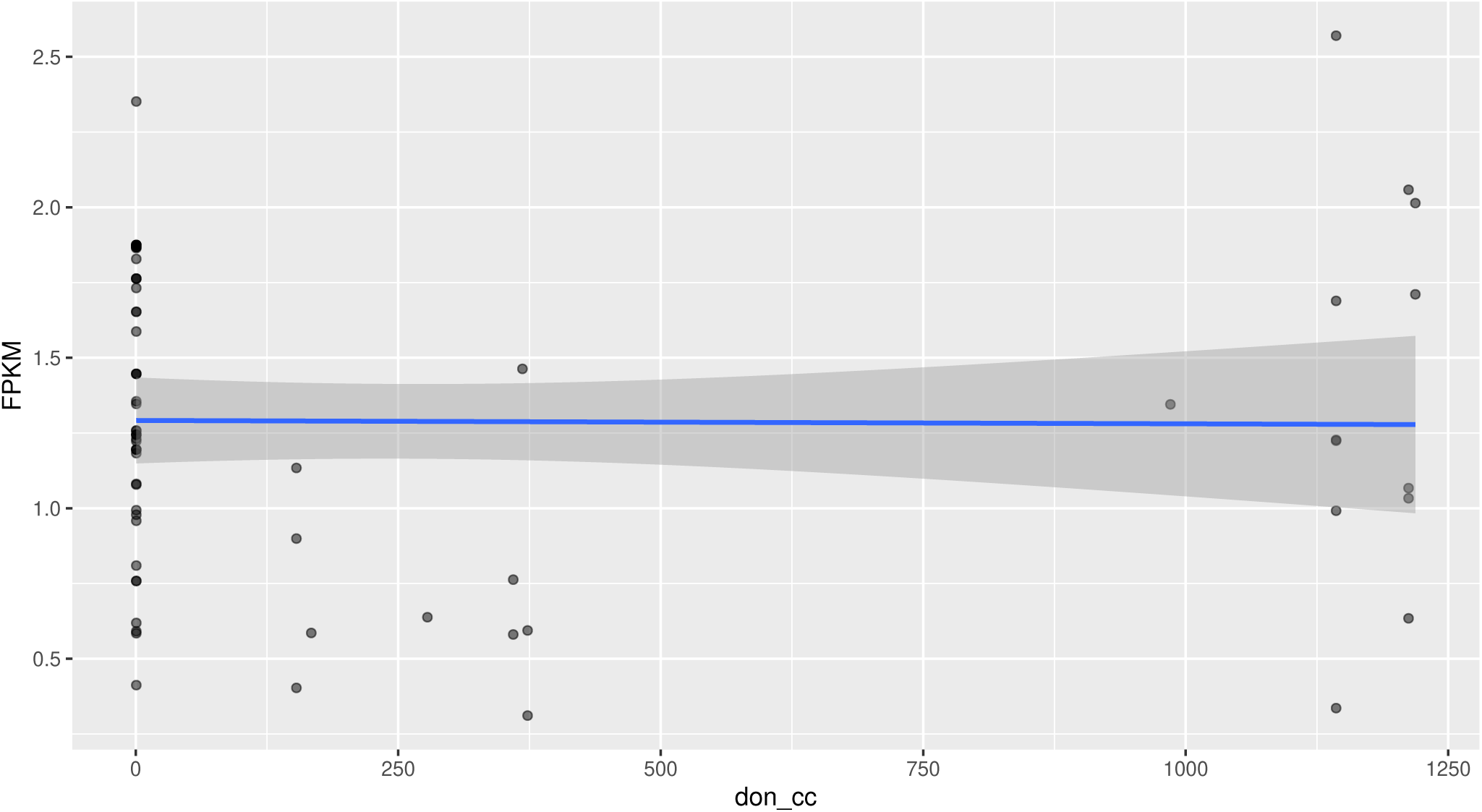
Relationship between DON concentration (don_cc) and ARG counts (FPKM). The scatter plot shows individual data points for ARG FPKM values across a range of DON concentrations, with a fitted linear regression line (blue).

**Figure 10.**
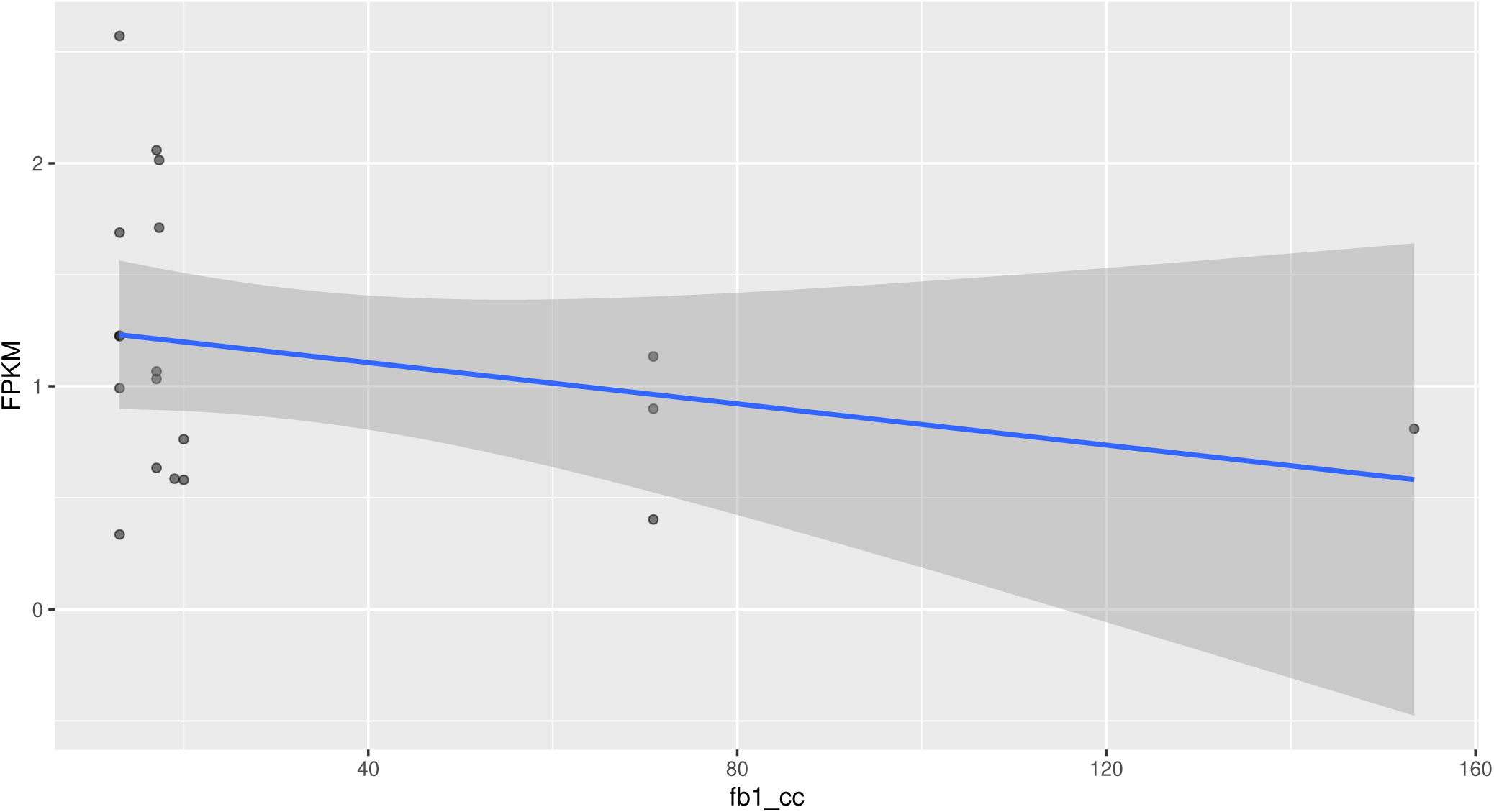
Relationship between FB1 concentration (fb1_cc) and ARG counts (FPKM). The scatter plot shows individual data points for ARG FPKM values across a range of FB1 concentrations, with a fitted linear regression line (blue).

**Figure 11.**
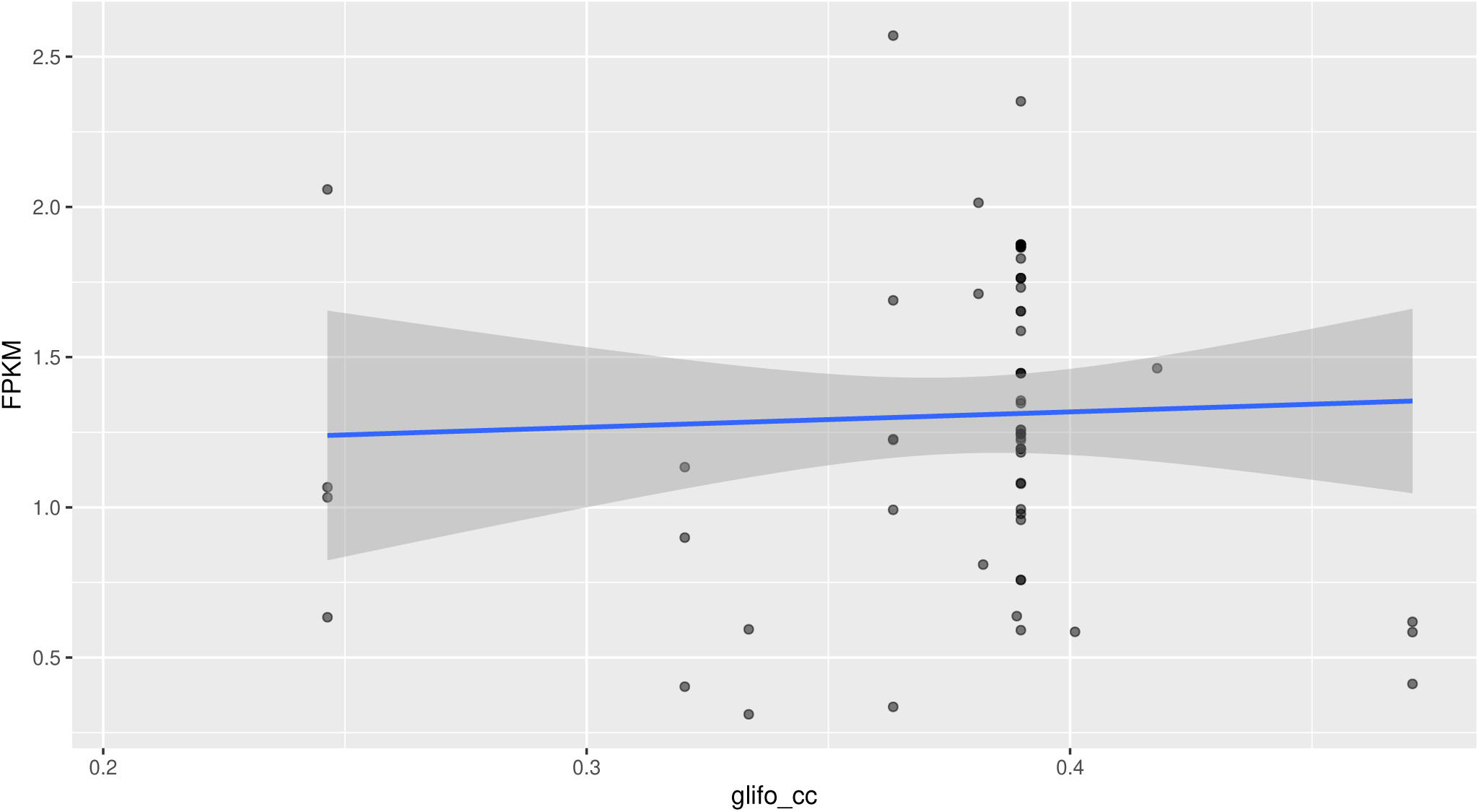
Relationship between glyphosate concentration (glifo_cc) and ARG counts (FPKM). The scatter plot shows individual data points for ARG FPKM values across a range of glyphosate concentrations, with a fitted linear regression line (blue).

**Figure 12.**
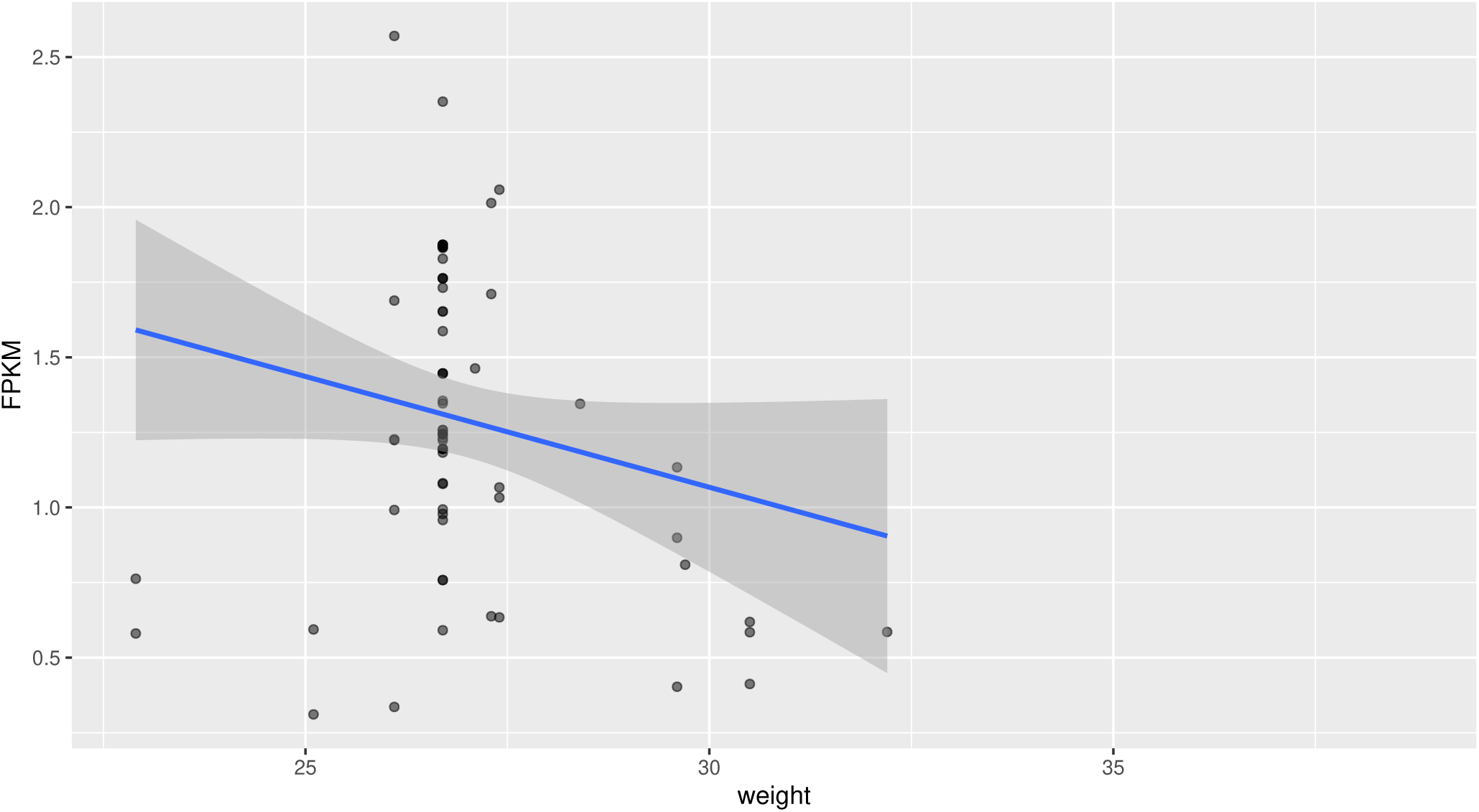
Relationship between weight (weight) and ARG counts (FPKM). The scatter plot shows individual data points for ARG FPKM values across a range of weight, with a fitted linear regression line (blue).

**Figure 13.**
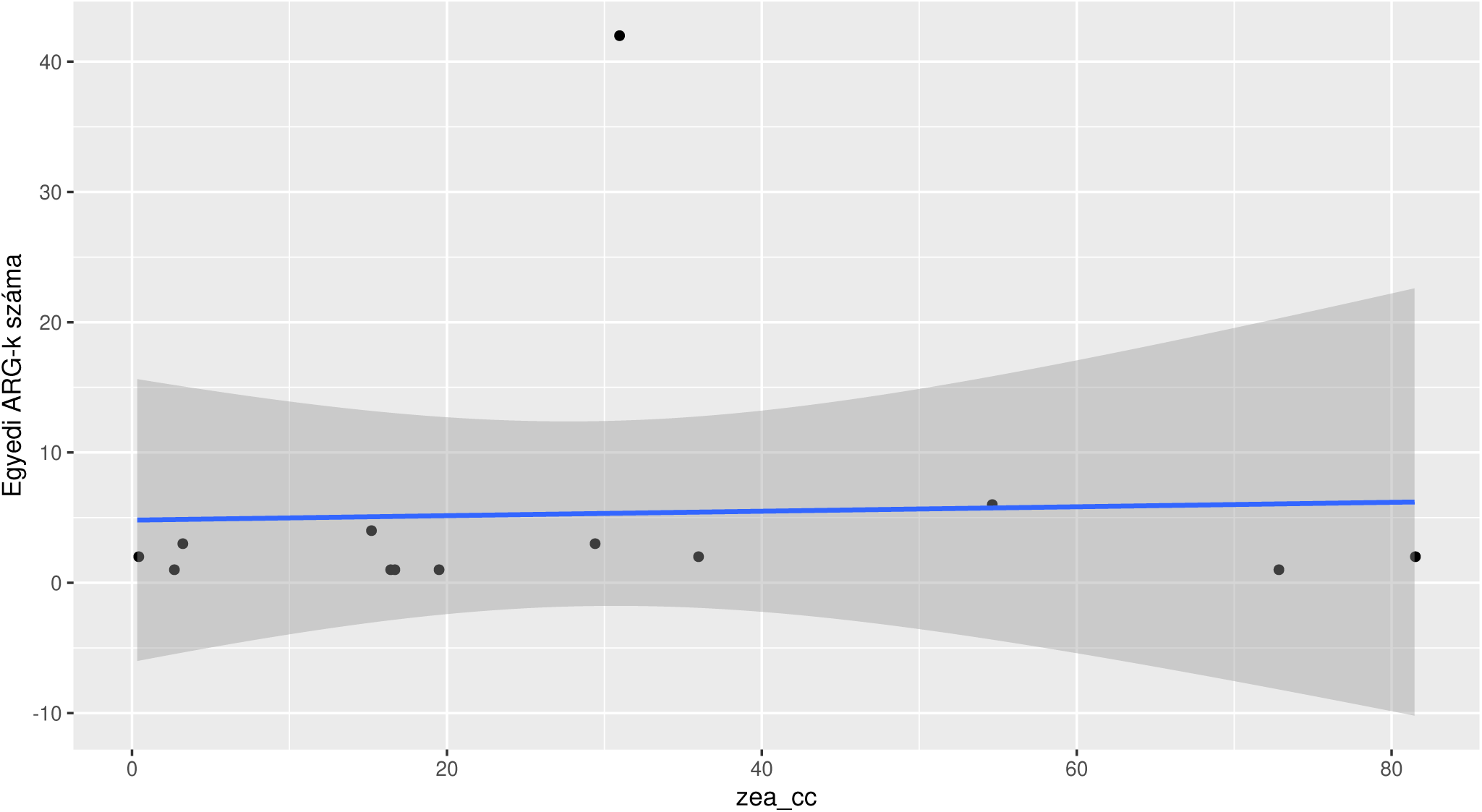
Relationship between ZEA concentration (zea_cc) and ARG counts. The scatter plot shows individual data points for ARG count values across a range of ZEA concentrations, with a fitted linear regression line (blue).

**Figure 14.**
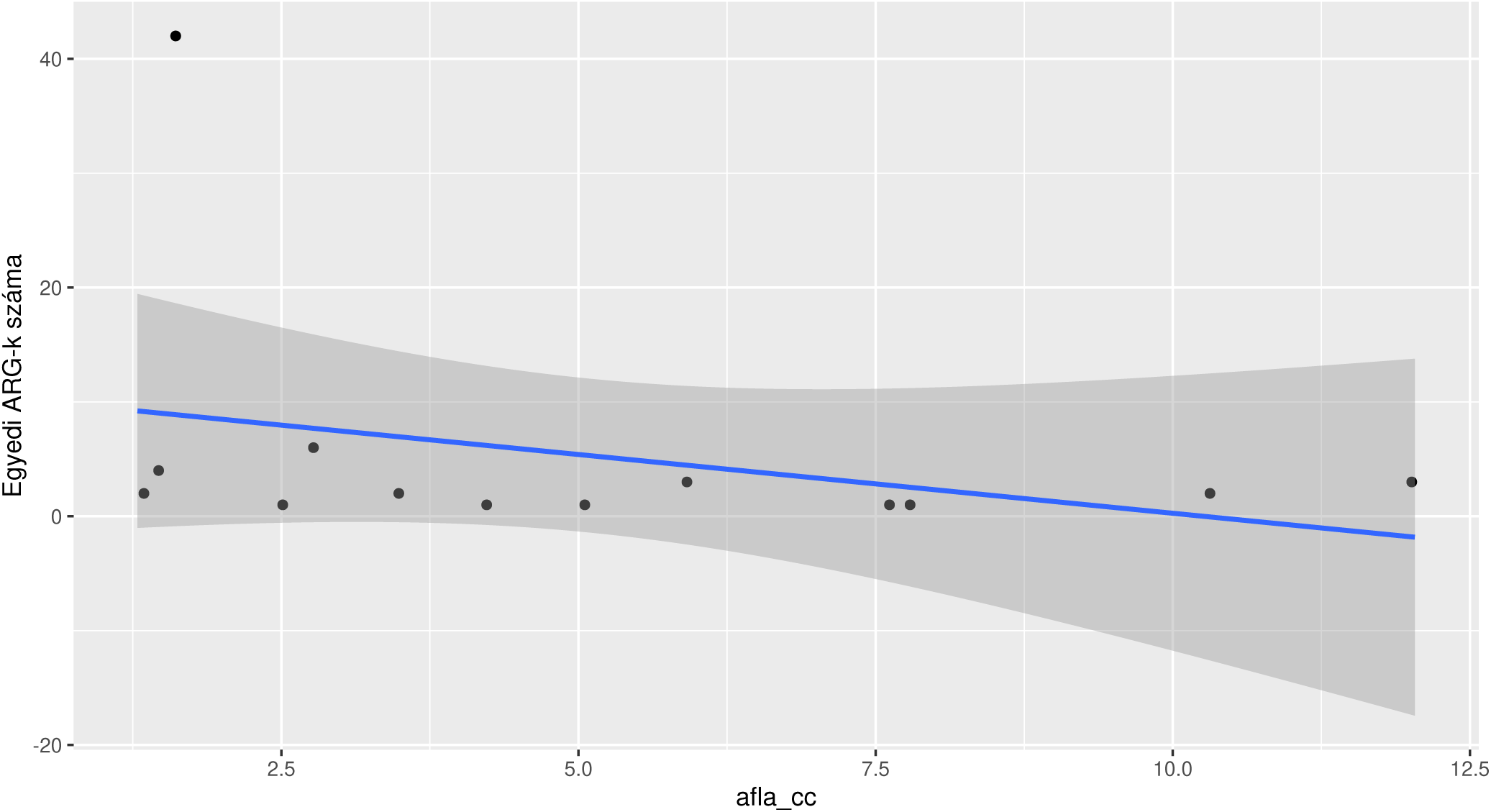
Relationship between AFs concentration (afla_cc) and ARG counts. The scatter plot shows individual data points for ARG count values across a range of AFs concentrations, with a fitted linear regression line (blue).

**Figure 15.**
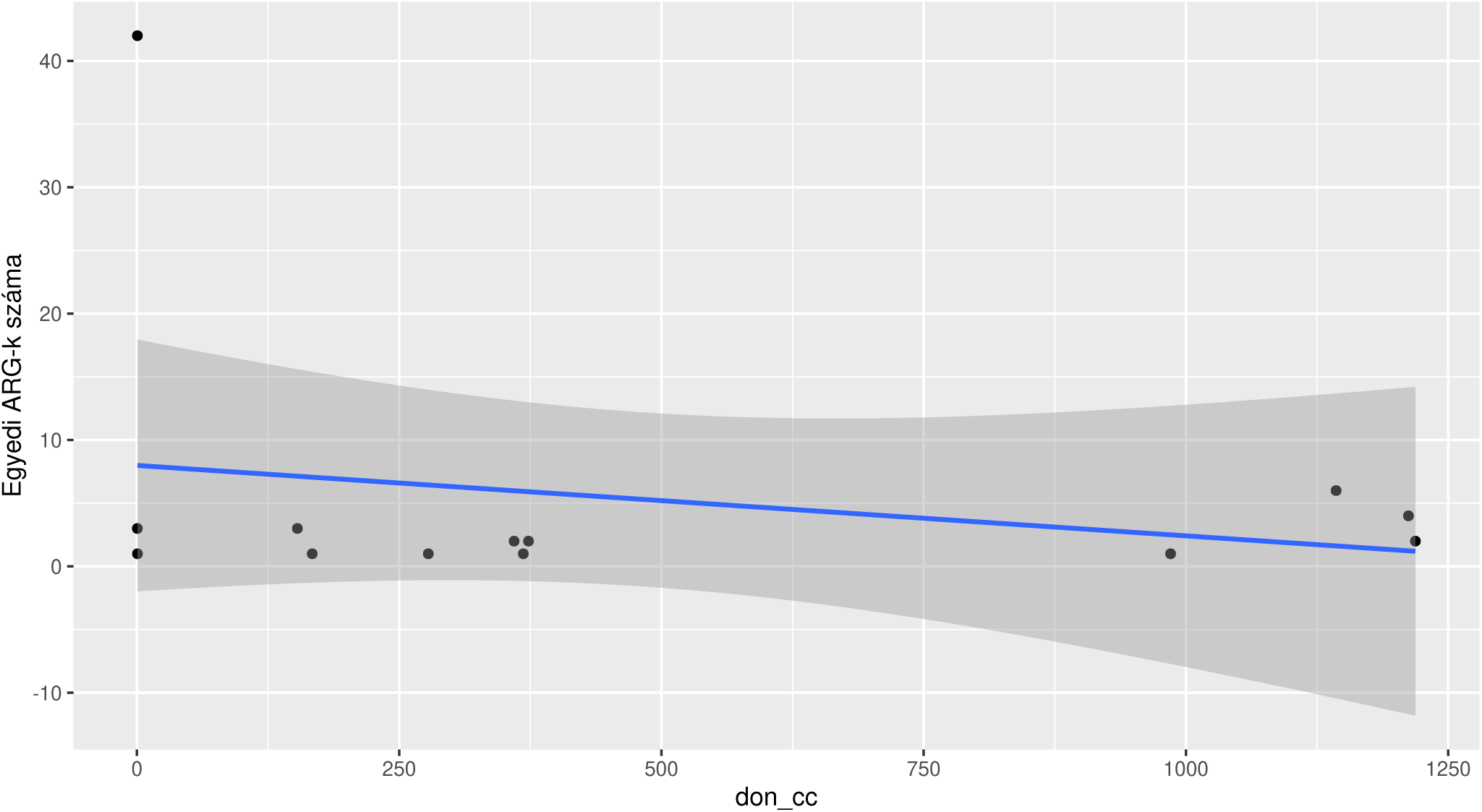
Relationship between DON concentration (don_cc) and ARG counts. The scatter plot shows individual data points for ARG count values across a range of DON concentrations, with a fitted linear regression line (blue).

**Figure 16.**
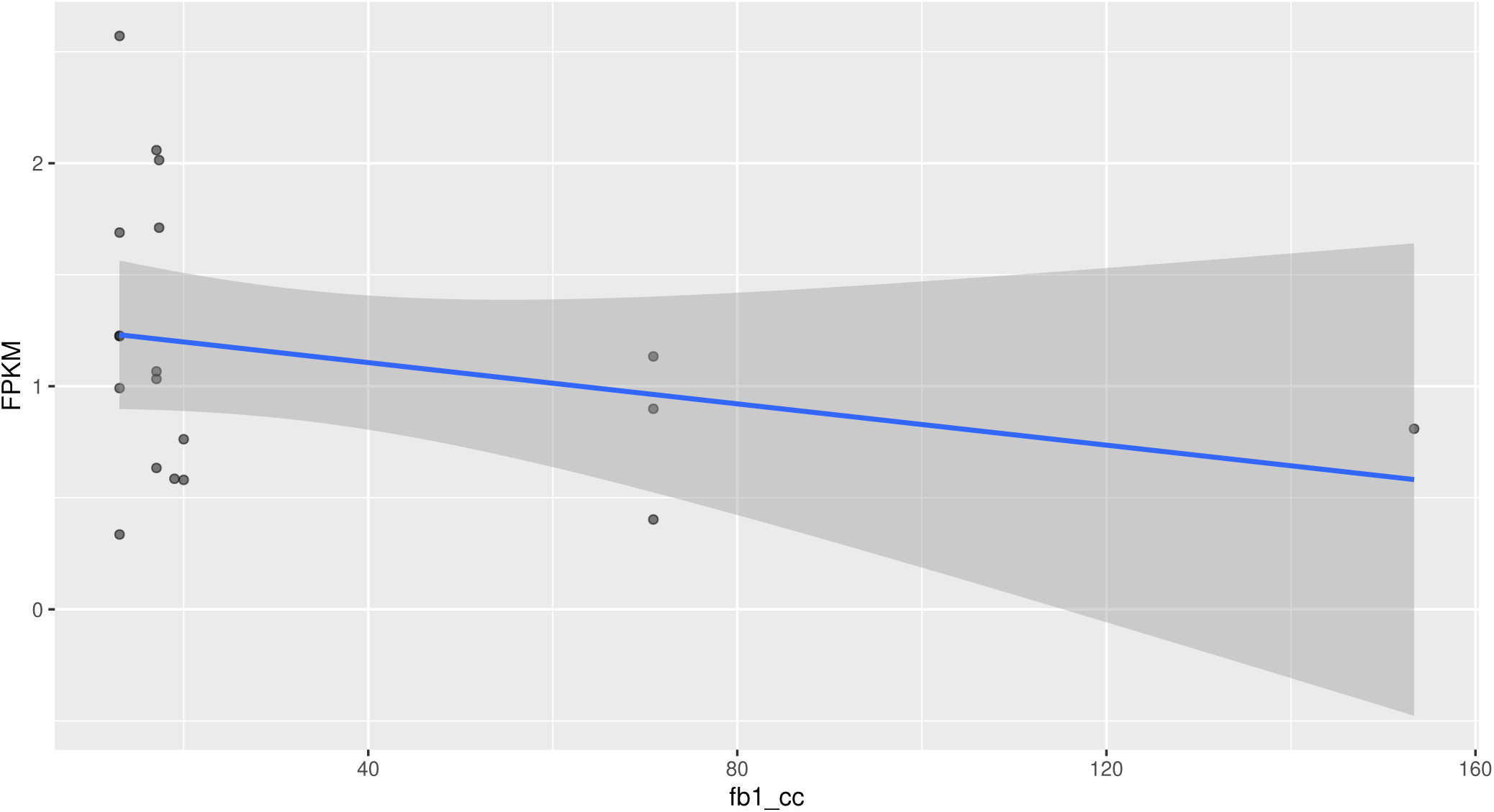
Relationship between FB1 concentration (fb1_cc) and ARG counts. The scatter plot shows individual data points for ARG count values across a range of FB1 concentrations, with a fitted linear regression line (blue).

**Figure 17.**
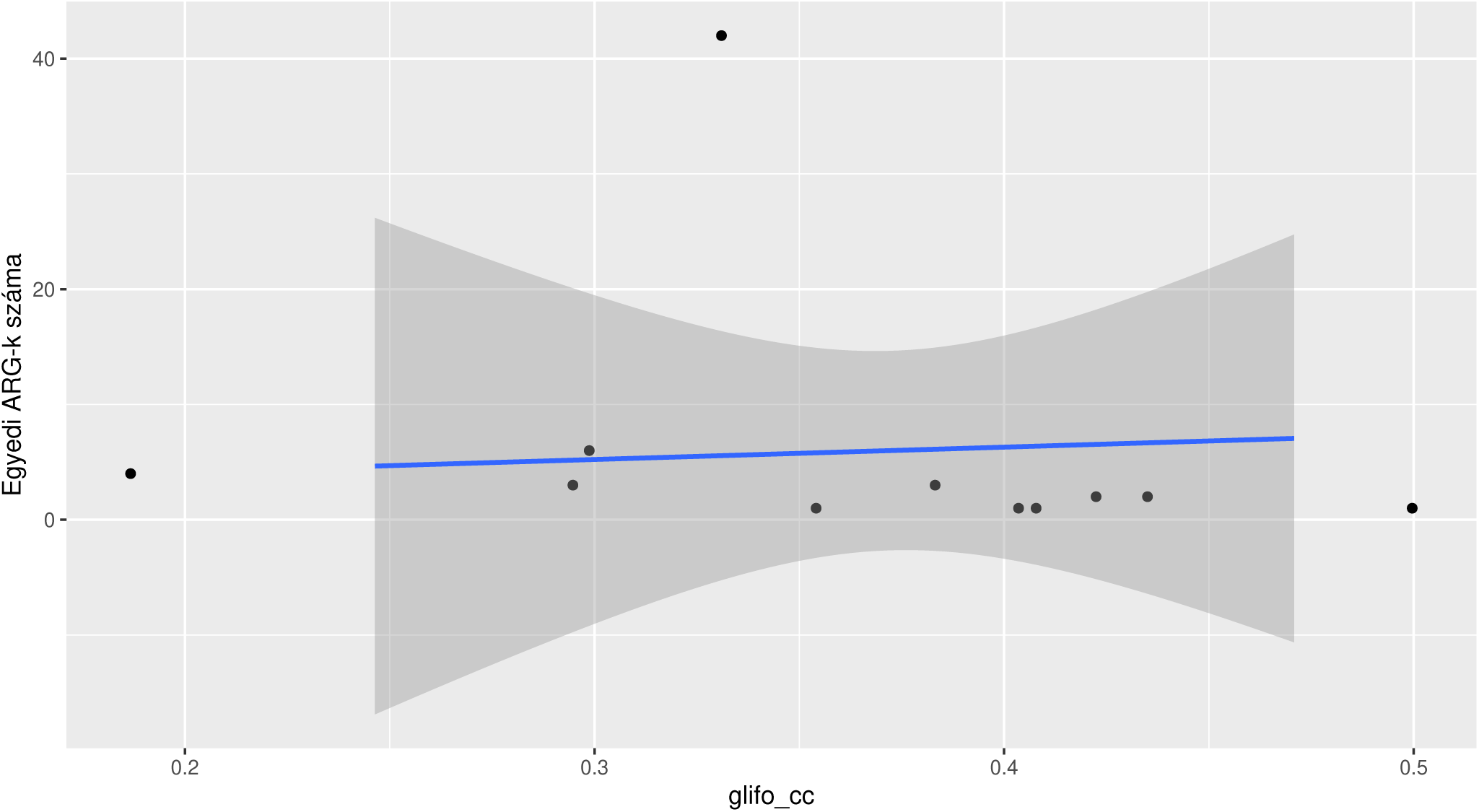
Relationship between glyphosate concentration (glifo_cc) and ARG counts. The scatter plot shows individual data points for ARG count values across a range of glyphosate concentrations, with a fitted linear regression line (blue).

**Figure 18.**
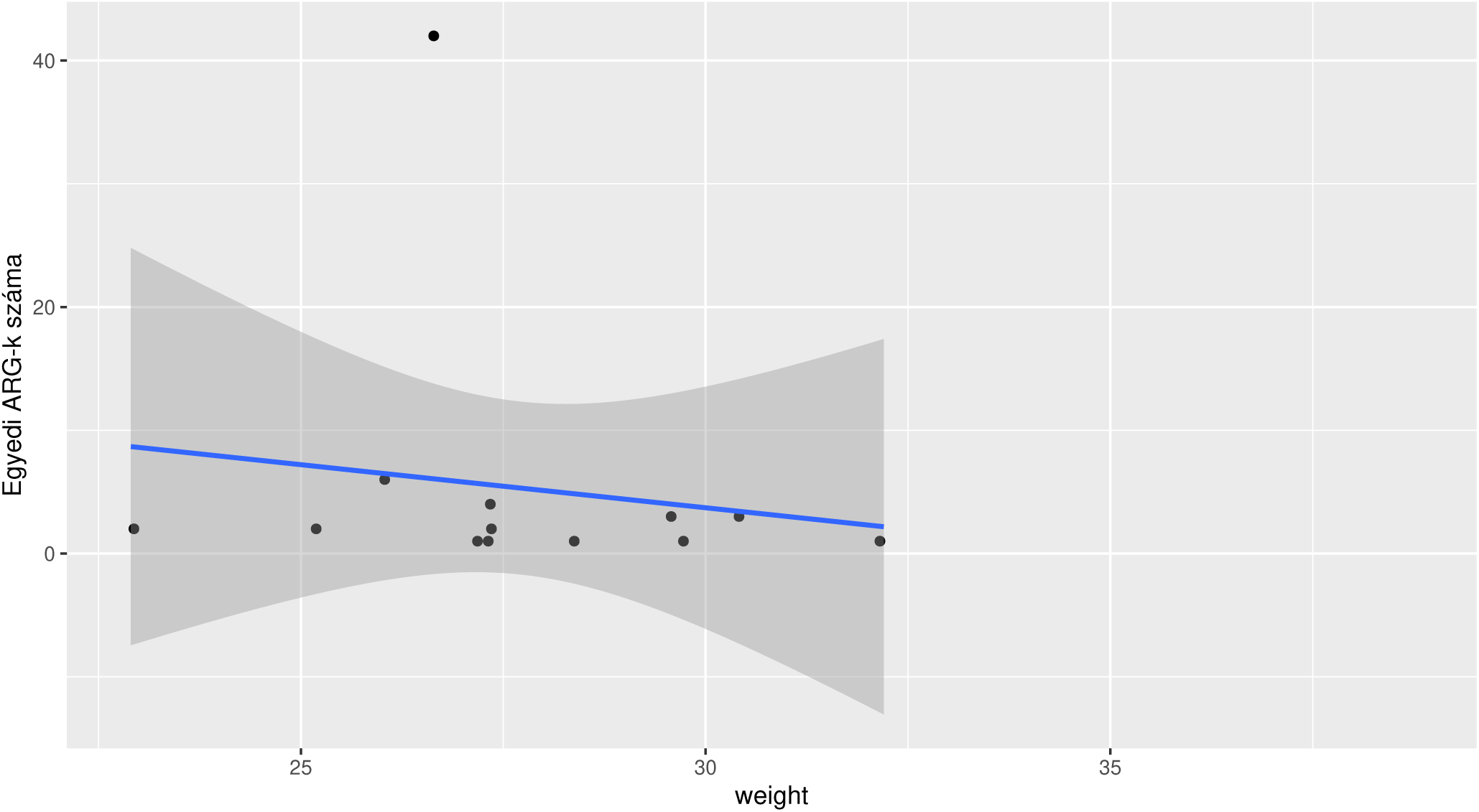
Relationship between weight (weight) and ARG counts. The scatter plot shows individual data points for ARG count values across a range of weight, with a fitted linear regression line (blue).

**Table 7.**
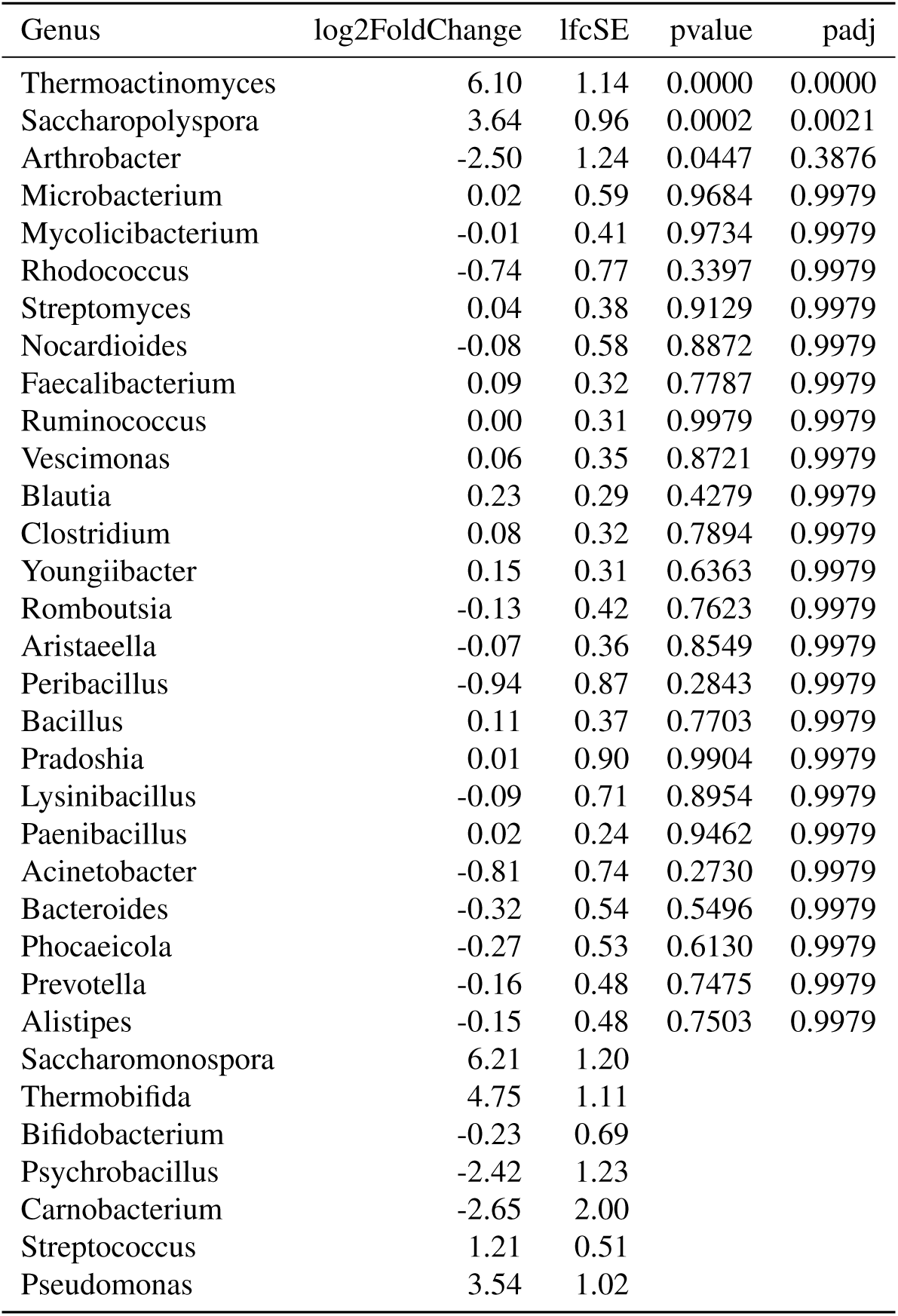
Differential abundance of microbial genera in response to varying ZEA levels. Genera are ranked by adjusted p-value (padj), with positive log2 fold change values indicating increased abundance at low ZEA level individuals compared to medium level ones.

**Table 8.** Differential abundance of microbial genera in response to varying ZEA levels. Genera are ranked by adjusted p-value (padj), with positive log2 fold change values indicating increased abundance at low ZEA level individuals compared to high level ones.

| Genus | log2FoldChange | lfcSE | pvalue | padj |
| --- | --- | --- | --- | --- |
| Thermoactinomyces | 5.92 | 1.14 | 0.0000 | 0.0000 |
| Rhodococcus | -3.37 | 0.77 | 0.0000 | 0.0002 |
| Saccharopolyspora | 3.54 | 0.96 | 0.0002 | 0.0020 |
| Acinetobacter | -2.68 | 0.74 | 0.0003 | 0.0020 |
| Nocardioides | -1.80 | 0.58 | 0.0018 | 0.0096 |
| Microbacterium | -1.42 | 0.59 | 0.0164 | 0.0709 |
| Romboutsia | -0.95 | 0.42 | 0.0248 | 0.0922 |
| Peribacillus | -1.69 | 0.87 | 0.0525 | 0.1708 |
| Mycolicibacterium | -0.76 | 0.41 | 0.0670 | 0.1867 |
| Aristaeella | -0.65 | 0.36 | 0.0718 | 0.1867 |
| Blautia | 0.49 | 0.29 | 0.0959 | 0.2266 |
| Arthrobacter | -1.99 | 1.24 | 0.1101 | 0.2385 |
| Faecalibacterium | 0.38 | 0.32 | 0.2281 | 0.4198 |
| Pradoshia | -1.11 | 0.90 | 0.2167 | 0.4198 |
| Lysinibacillus | -0.83 | 0.71 | 0.2462 | 0.4198 |
| Alistipes | 0.54 | 0.48 | 0.2583 | 0.4198 |
| Vescimonas | 0.30 | 0.35 | 0.3947 | 0.6037 |
| Clostridium | 0.23 | 0.32 | 0.4748 | 0.6497 |
| Phocaeicola | 0.40 | 0.53 | 0.4536 | 0.6497 |
| Bacillus | -0.23 | 0.37 | 0.5218 | 0.6783 |
| Streptomyces | -0.18 | 0.38 | 0.6303 | 0.7216 |
| Ruminococcus | -0.17 | 0.31 | 0.5834 | 0.7216 |
| Bacteroides | 0.25 | 0.54 | 0.6383 | 0.7216 |
| Prevotella | 0.18 | 0.48 | 0.7103 | 0.7695 |
| Paenibacillus | -0.05 | 0.24 | 0.8451 | 0.8789 |
| Youngiibacter | -0.00 | 0.31 | 0.9904 | 0.9904 |
| Saccharomonospora | 6.28 | 1.20 |  |  |
| Thermobifida | 4.68 | 1.11 |  |  |
| Bifidobacterium | -1.57 | 0.69 |  |  |
| Psychrobacillus | -1.26 | 1.23 |  |  |
| Carnobacterium | -1.91 | 2.00 |  |  |
| Streptococcus | 1.19 | 0.51 |  |  |
| Pseudomonas | 3.98 | 1.02 |  |  |

**Table 9.** Differential abundance of microbial genera in response to varying ZEA levels. Genera are ranked by adjusted p-value (padj), with positive log2 fold change values indicating increased abundance at medium ZEA level individuals compared to high level ones.

| Genus | log2FoldChange | lfcSE | pvalue | padj |
| --- | --- | --- | --- | --- |
| Rhodococcus | -2.63 | 0.77 | 0.0007 | 0.0177 |
| Nocardioides | -1.72 | 0.58 | 0.0029 | 0.0383 |
| Microbacterium | -1.44 | 0.59 | 0.0147 | 0.0954 |
| Acinetobacter | -1.87 | 0.74 | 0.0119 | 0.0954 |
| Romboutsia | -0.82 | 0.42 | 0.0522 | 0.2713 |
| Mycolicibacterium | -0.74 | 0.41 | 0.0721 | 0.3124 |
| Aristaeella | -0.58 | 0.36 | 0.1058 | 0.3929 |
| Alistipes | 0.69 | 0.48 | 0.1474 | 0.4792 |
| Pradoshia | -1.12 | 0.90 | 0.2123 | 0.5519 |
| Phocaeicola | 0.67 | 0.53 | 0.2094 | 0.5519 |
| Faecalibacterium | 0.29 | 0.32 | 0.3554 | 0.6263 |
| Blautia | 0.26 | 0.29 | 0.3830 | 0.6263 |
| Peribacillus | -0.76 | 0.87 | 0.3854 | 0.6263 |
| Bacillus | -0.34 | 0.37 | 0.3510 | 0.6263 |
| Lysinibacillus | -0.73 | 0.71 | 0.3039 | 0.6263 |
| Bacteroides | 0.58 | 0.54 | 0.2853 | 0.6263 |
| Vescimonas | 0.24 | 0.35 | 0.4901 | 0.7080 |
| Prevotella | 0.34 | 0.48 | 0.4881 | 0.7080 |
| Streptomyces | -0.23 | 0.38 | 0.5547 | 0.7561 |
| Ruminococcus | -0.17 | 0.31 | 0.5816 | 0.7561 |
| Arthrobacter | 0.51 | 1.24 | 0.6822 | 0.7712 |
| Clostridium | 0.14 | 0.32 | 0.6544 | 0.7712 |
| Youngiibacter | -0.15 | 0.31 | 0.6277 | 0.7712 |
| Paenibacillus | -0.06 | 0.24 | 0.7927 | 0.8587 |
| Thermoactinomyces | -0.18 | 1.15 | 0.8740 | 0.9089 |
| Saccharopolyspora | -0.10 | 0.96 | 0.9160 | 0.9160 |
| Saccharomonospora | 0.08 | 1.20 |  |  |
| Thermobifida | -0.07 | 1.11 |  |  |
| Bifidobacterium | -1.33 | 0.69 |  |  |
| Psychrobacillus | 1.16 | 1.23 |  |  |
| Carnobacterium | 0.74 | 2.00 |  |  |
| Streptococcus | -0.02 | 0.51 |  |  |
| Pseudomonas | 0.44 | 1.02 |  |  |

**Table 10.** Differential abundance of microbial genera in response to varying ZEA concentrations. Genera are ranked by adjusted p-value (padj), with positive log2 fold change values indicating increased abundance at higher ZEA concentrations.

| Genus | log2FoldChange | lfcSE | pvalue | padj |
| --- | --- | --- | --- | --- |
| Rhodococcus | 0.06 | 0.02 | 0.0001 | 0.0018 |
| Acinetobacter | 0.05 | 0.01 | 0.0006 | 0.0107 |
| Bifidobacterium | 0.03 | 0.01 | 0.0010 | 0.0112 |
| Nocardioides | 0.03 | 0.01 | 0.0033 | 0.0272 |
| Romboutsia | 0.02 | 0.01 | 0.0057 | 0.0374 |
| Thermobifida | -0.06 | 0.02 | 0.0130 | 0.0535 |
| Aristaeella | 0.01 | 0.01 | 0.0128 | 0.0535 |
| Pseudomonas | -0.05 | 0.02 | 0.0101 | 0.0535 |
| Saccharopolyspora | -0.04 | 0.02 | 0.0258 | 0.0944 |
| Microbacterium | 0.02 | 0.01 | 0.0384 | 0.1268 |
| Mycolicibacterium | 0.01 | 0.01 | 0.0479 | 0.1437 |
| Peribacillus | 0.03 | 0.01 | 0.0529 | 0.1453 |
| Streptococcus | -0.01 | 0.01 | 0.1278 | 0.3245 |
| Blautia | -0.01 | 0.00 | 0.2103 | 0.4958 |
| Alistipes | -0.01 | 0.01 | 0.3356 | 0.7384 |
| Streptomyces | 0.00 | 0.01 | 0.4048 | 0.8060 |
| Faecalibacterium | -0.00 | 0.01 | 0.4396 | 0.8060 |
| Ruminococcus | 0.00 | 0.00 | 0.4209 | 0.8060 |
| Arthrobacter | 0.01 | 0.02 | 0.6281 | 0.8342 |
| Saccharomonospora | -0.02 | 0.06 | 0.7331 | 0.8342 |
| Vescimonas | -0.00 | 0.01 | 0.6229 | 0.8342 |
| Youngiibacter | 0.00 | 0.00 | 0.6191 | 0.8342 |
| Psychrobacillus | 0.01 | 0.02 | 0.7043 | 0.8342 |
| Pradoshia | 0.01 | 0.02 | 0.5492 | 0.8342 |
| Thermoactinomyces | -0.02 | 0.06 | 0.7072 | 0.8342 |
| Carnobacterium | 0.02 | 0.06 | 0.7274 | 0.8342 |
| Bacteroides | -0.00 | 0.01 | 0.7140 | 0.8342 |
| Phocaeicola | -0.01 | 0.01 | 0.5236 | 0.8342 |
| Prevotella | -0.00 | 0.01 | 0.6741 | 0.8342 |
| Lysinibacillus | 0.00 | 0.01 | 0.7710 | 0.8481 |
| Bacillus | 0.00 | 0.01 | 0.8147 | 0.8611 |
| Paenibacillus | 0.00 | 0.00 | 0.8350 | 0.8611 |
| Clostridium | -0.00 | 0.00 | 0.9278 | 0.9278 |

**Table 11.**
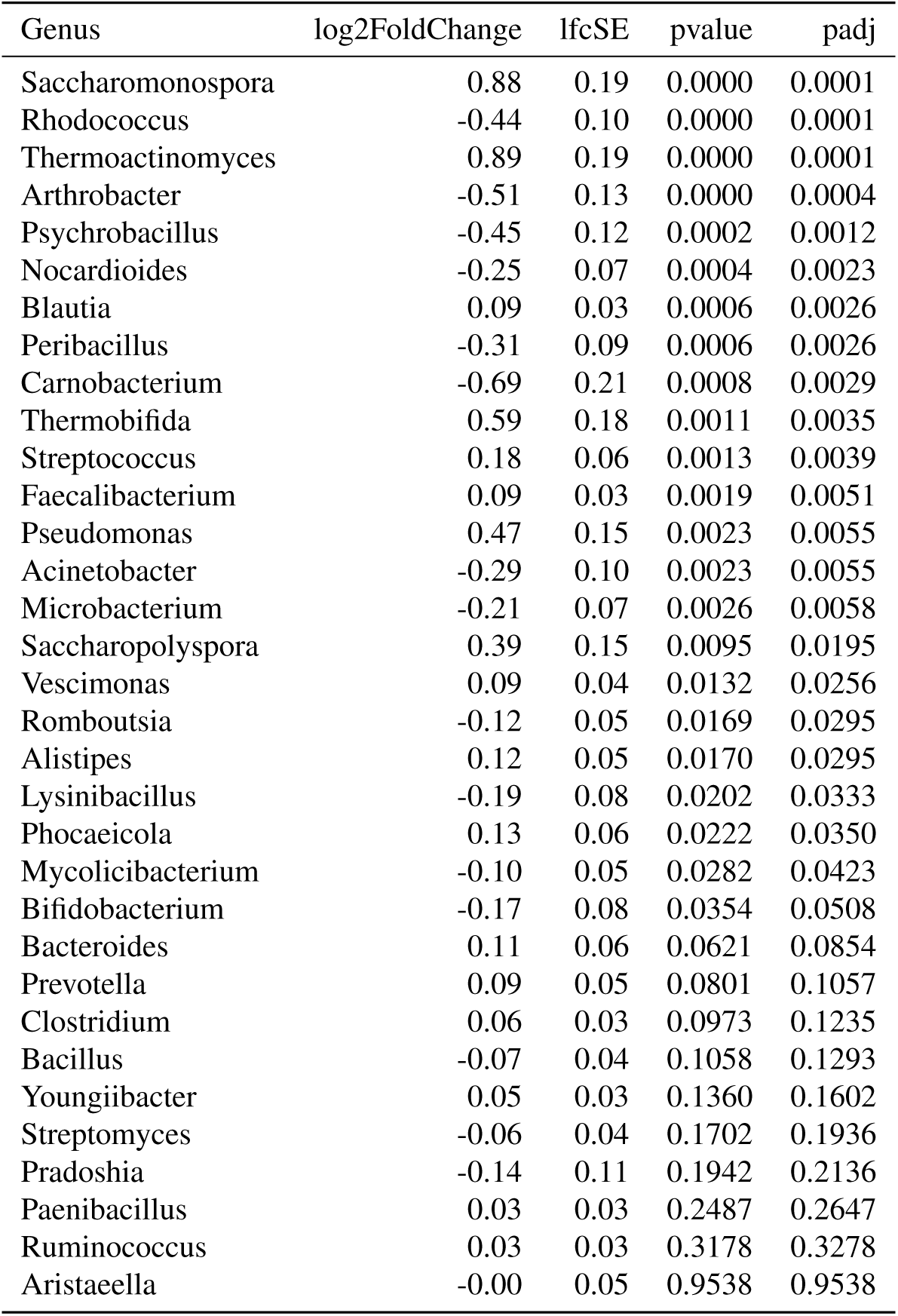
Differential abundance of microbial genera in response to varying AFs concentrations. Genera are ranked by adjusted p-value (padj), with positive log2 fold change values indicating increased abundance at higher AFs concentrations.

| Genus | log2FoldChange | lfcSE | pvalue | padj |
| --- | --- | --- | --- | --- |
| Saccharomonospora | 0.88 | 0.19 | 0.0000 | 0.0001 |
| Rhodococcus | -0.44 | 0.10 | 0.0000 | 0.0001 |
| Thermoactinomyces | 0.89 | 0.19 | 0.0000 | 0.0001 |
| Arthrobacter | -0.51 | 0.13 | 0.0000 | 0.0004 |
| Psychrobacillus | -0.45 | 0.12 | 0.0002 | 0.0012 |
| Nocardioides | -0.25 | 0.07 | 0.0004 | 0.0023 |
| Blautia | 0.09 | 0.03 | 0.0006 | 0.0026 |
| Peribacillus | -0.31 | 0.09 | 0.0006 | 0.0026 |
| Carnobacterium | -0.69 | 0.21 | 0.0008 | 0.0029 |
| Thermobifida | 0.59 | 0.18 | 0.0011 | 0.0035 |
| Streptococcus | 0.18 | 0.06 | 0.0013 | 0.0039 |
| Faecalibacterium | 0.09 | 0.03 | 0.0019 | 0.0051 |
| Pseudomonas | 0.47 | 0.15 | 0.0023 | 0.0055 |
| Acinetobacter | -0.29 | 0.10 | 0.0023 | 0.0055 |
| Microbacterium | -0.21 | 0.07 | 0.0026 | 0.0058 |
| Saccharopolyspora | 0.39 | 0.15 | 0.0095 | 0.0195 |
| Vescimonas | 0.09 | 0.04 | 0.0132 | 0.0256 |
| Romboutsia | -0.12 | 0.05 | 0.0169 | 0.0295 |
| Alistipes | 0.12 | 0.05 | 0.0170 | 0.0295 |
| Lysinibacillus | -0.19 | 0.08 | 0.0202 | 0.0333 |
| Phocaeicola | 0.13 | 0.06 | 0.0222 | 0.0350 |
| Mycolicibacterium | -0.10 | 0.05 | 0.0282 | 0.0423 |
| Bifidobacterium | -0.17 | 0.08 | 0.0354 | 0.0508 |
| Bacteroides | 0.11 | 0.06 | 0.0621 | 0.0854 |
| Prevotella | 0.09 | 0.05 | 0.0801 | 0.1057 |
| Clostridium | 0.06 | 0.03 | 0.0973 | 0.1235 |
| Bacillus | -0.07 | 0.04 | 0.1058 | 0.1293 |
| Youngiibacter | 0.05 | 0.03 | 0.1360 | 0.1602 |
| Streptomyces | -0.06 | 0.04 | 0.1702 | 0.1936 |
| Pradoshia | -0.14 | 0.11 | 0.1942 | 0.2136 |
| Paenibacillus | 0.03 | 0.03 | 0.2487 | 0.2647 |
| Ruminococcus | 0.03 | 0.03 | 0.3178 | 0.3278 |
| Aristaeella | -0.00 | 0.05 | 0.9538 | 0.9538 |

**Table 12.**
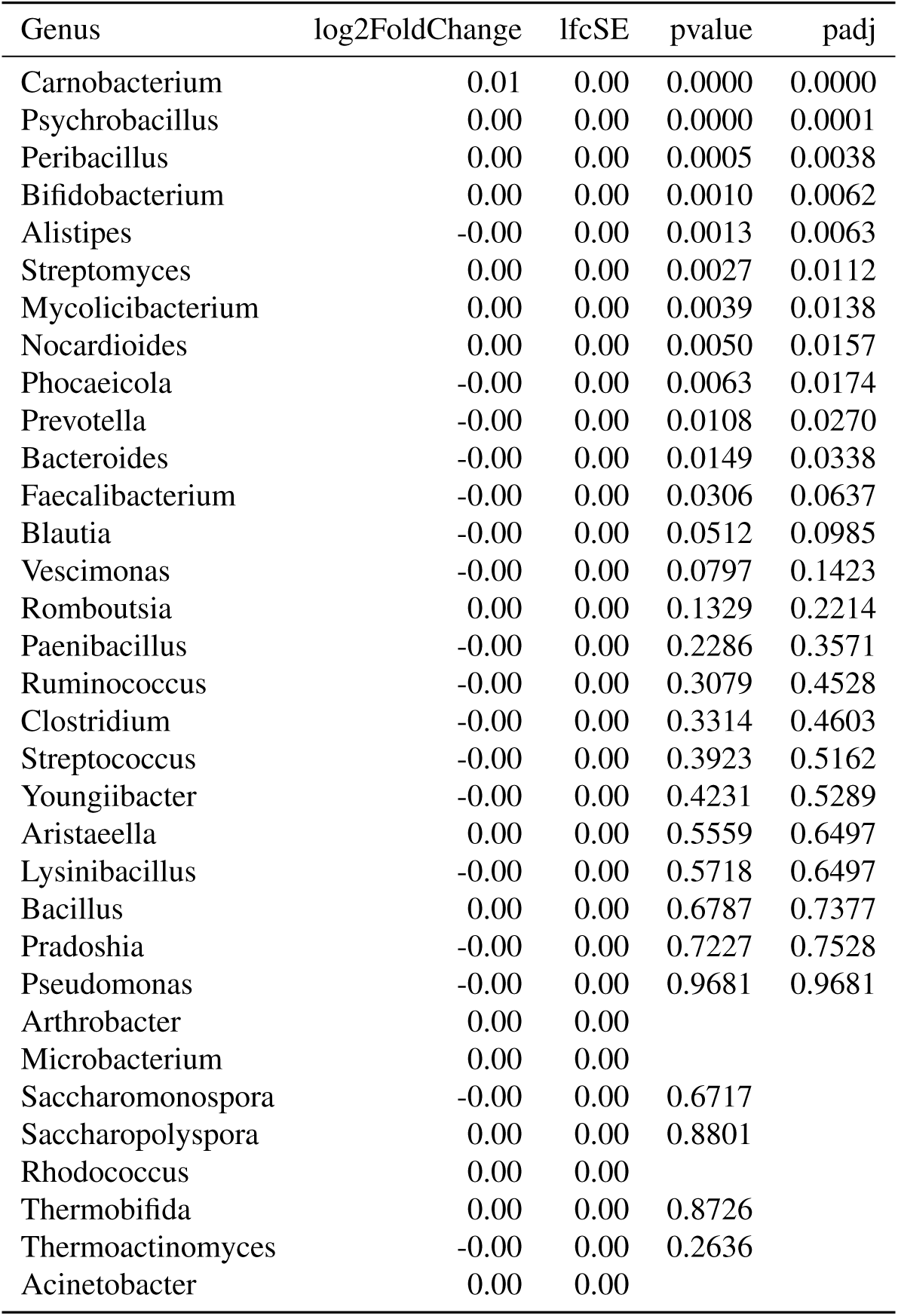
Differential abundance of microbial genera in response to varying DON concentrations. Genera are ranked by adjusted p-value (padj), with positive log2 fold change values indicating increased abundance at higher DON concentrations.

| Genus | log2FoldChange | lfcSE | pvalue | padj |
| --- | --- | --- | --- | --- |
| Carnobacterium | 0.01 | 0.00 | 0.0000 | 0.0000 |
| Psychrobacillus | 0.00 | 0.00 | 0.0000 | 0.0001 |
| Peribacillus | 0.00 | 0.00 | 0.0005 | 0.0038 |
| Bifidobacterium | 0.00 | 0.00 | 0.0010 | 0.0062 |
| Alistipes | -0.00 | 0.00 | 0.0013 | 0.0063 |
| Streptomyces | 0.00 | 0.00 | 0.0027 | 0.0112 |
| Mycolicibacterium | 0.00 | 0.00 | 0.0039 | 0.0138 |
| Nocardioides | 0.00 | 0.00 | 0.0050 | 0.0157 |
| Phocaeicola | -0.00 | 0.00 | 0.0063 | 0.0174 |
| Prevotella | -0.00 | 0.00 | 0.0108 | 0.0270 |
| Bacteroides | -0.00 | 0.00 | 0.0149 | 0.0338 |
| Faecalibacterium | -0.00 | 0.00 | 0.0306 | 0.0637 |
| Blautia | -0.00 | 0.00 | 0.0512 | 0.0985 |
| Vescimonas | -0.00 | 0.00 | 0.0797 | 0.1423 |
| Romboutsia | 0.00 | 0.00 | 0.1329 | 0.2214 |
| Paenibacillus | -0.00 | 0.00 | 0.2286 | 0.3571 |
| Ruminococcus | -0.00 | 0.00 | 0.3079 | 0.4528 |
| Clostridium | -0.00 | 0.00 | 0.3314 | 0.4603 |
| Streptococcus | -0.00 | 0.00 | 0.3923 | 0.5162 |
| Youngiibacter | -0.00 | 0.00 | 0.4231 | 0.5289 |
| Aristaeella | 0.00 | 0.00 | 0.5559 | 0.6497 |
| Lysinibacillus | -0.00 | 0.00 | 0.5718 | 0.6497 |
| Bacillus | 0.00 | 0.00 | 0.6787 | 0.7377 |
| Pradoshia | -0.00 | 0.00 | 0.7227 | 0.7528 |
| Pseudomonas | -0.00 | 0.00 | 0.9681 | 0.9681 |
| Arthrobacter | 0.00 | 0.00 |  |  |
| Microbacterium | 0.00 | 0.00 |  |  |
| Saccharomonospora | -0.00 | 0.00 | 0.6717 |  |
| Saccharopolyspora | 0.00 | 0.00 | 0.8801 |  |
| Rhodococcus | 0.00 | 0.00 |  |  |
| Thermobifida | 0.00 | 0.00 | 0.8726 |  |
| Thermoactinomyces | -0.00 | 0.00 | 0.2636 |  |
| Acinetobacter | 0.00 | 0.00 |  |  |

**Table 13.**
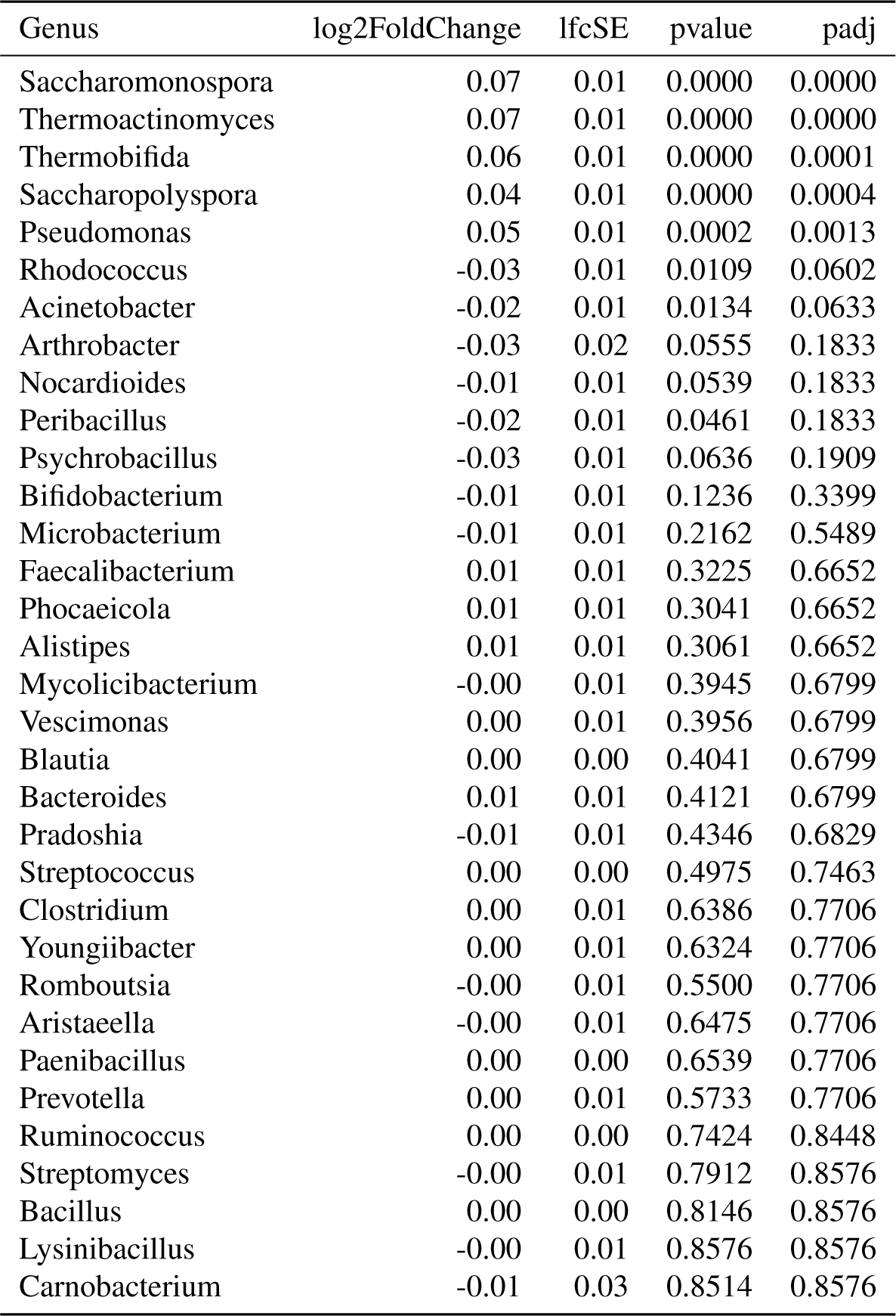
Differential abundance of microbial genera in response to varying FB1 concentrations. Genera are ranked by adjusted p-value (padj), with positive log2 fold change values indicating increased abundance at higher FB1 concentrations.

**Table 14.** Differential abundance of microbial genera in response to varying glyphosate concentrations. Genera are ranked by adjusted p-value (padj), with positive log2 fold change values indicating increased abundance at higher glyphosate concentrations.

| Genus | log2FoldChange | lfcSE | pvalue | padj |
| --- | --- | --- | --- | --- |
| Saccharomonospora | -29.87 | 8.78 | 0.0007 | 0.0055 |
| Thermobifida | -27.35 | 7.31 | 0.0002 | 0.0055 |
| Thermoactinomyces | -30.00 | 8.39 | 0.0003 | 0.0055 |
| Pseudomonas | -22.56 | 6.57 | 0.0006 | 0.0055 |
| Saccharopolyspora | -21.18 | 6.38 | 0.0009 | 0.0060 |
| Psychrobacillus | -16.64 | 6.71 | 0.0131 | 0.0723 |
| Lysinibacillus | 9.00 | 3.94 | 0.0224 | 0.1057 |
| Pradoshia | 11.52 | 5.44 | 0.0341 | 0.1405 |
| Arthrobacter | -14.53 | 7.93 | 0.0669 | 0.1885 |
| Streptomyces | -3.95 | 2.14 | 0.0647 | 0.1885 |
| Romboutsia | 5.45 | 2.95 | 0.0642 | 0.1885 |
| Carnobacterium | -20.28 | 11.13 | 0.0685 | 0.1885 |
| Prevotella | 4.39 | 2.94 | 0.1361 | 0.3454 |
| Bacteroides | 4.53 | 3.25 | 0.1634 | 0.3851 |
| Alistipes | 4.19 | 3.10 | 0.1765 | 0.3884 |
| Phocaeicola | 4.28 | 3.38 | 0.2059 | 0.4247 |
| Mycolicibacterium | -2.91 | 2.93 | 0.3212 | 0.5888 |
| Ruminococcus | 1.68 | 1.67 | 0.3153 | 0.5888 |
| Paenibacillus | 1.31 | 1.41 | 0.3495 | 0.6071 |
| Microbacterium | -2.40 | 4.52 | 0.5946 | 0.8267 |
| Bifidobacterium | 2.72 | 5.05 | 0.5904 | 0.8267 |
| Faecalibacterium | 1.01 | 2.08 | 0.6263 | 0.8267 |
| Clostridium | 1.18 | 1.83 | 0.5179 | 0.8267 |
| Peribacillus | -3.17 | 6.16 | 0.6070 | 0.8267 |
| Streptococcus | -2.20 | 3.77 | 0.5598 | 0.8267 |
| Vescimonas | 0.99 | 2.22 | 0.6542 | 0.8303 |
| Rhodococcus | 2.12 | 6.97 | 0.7607 | 0.8528 |
| Blautia | 0.50 | 1.91 | 0.7949 | 0.8528 |
| Youngiibacter | 0.40 | 1.81 | 0.8250 | 0.8528 |
| Aristaeella | 0.52 | 2.36 | 0.8270 | 0.8528 |
| Bacillus | 0.50 | 2.24 | 0.8225 | 0.8528 |
| Acinetobacter | 1.85 | 5.99 | 0.7571 | 0.8528 |
| Nocardioides | -0.43 | 4.91 | 0.9303 | 0.9303 |

**Table 15.** Differential abundance of microbial genera in response to varying weights. Genera are ranked by adjusted p-value (padj), with positive log2 fold change values indicating increased abundance at higher weights.

| Genus | log2FoldChange | lfcSE | pvalue | padj |
| --- | --- | --- | --- | --- |
| Pseudomonas | 0.51 | 0.13 | 0.0001 | 0.0017 |
| Thermobifida | 0.54 | 0.16 | 0.0006 | 0.0104 |
| Arthrobacter | -0.43 | 0.14 | 0.0016 | 0.0173 |
| Romboutsia | -0.14 | 0.05 | 0.0025 | 0.0206 |
| Rhodococcus | -0.30 | 0.11 | 0.0052 | 0.0346 |
| Psychrobacillus | -0.29 | 0.13 | 0.0292 | 0.1606 |
| Saccharopolyspora | 0.30 | 0.14 | 0.0351 | 0.1655 |
| Acinetobacter | -0.19 | 0.10 | 0.0674 | 0.2780 |
| Nocardioides | -0.14 | 0.08 | 0.0886 | 0.3249 |
| Microbacterium | -0.10 | 0.08 | 0.1794 | 0.4078 |
| Faecalibacterium | 0.05 | 0.03 | 0.1777 | 0.4078 |
| Vescimonas | 0.05 | 0.04 | 0.1582 | 0.4078 |
| Clostridium | 0.04 | 0.03 | 0.1948 | 0.4078 |
| Peribacillus | -0.14 | 0.11 | 0.1804 | 0.4078 |
| Pradoshia | -0.15 | 0.10 | 0.1273 | 0.4078 |
| Streptococcus | -0.08 | 0.07 | 0.1977 | 0.4078 |
| Blautia | 0.04 | 0.03 | 0.2240 | 0.4107 |
| Lysinibacillus | -0.10 | 0.08 | 0.2143 | 0.4107 |
| Alistipes | 0.06 | 0.06 | 0.2889 | 0.5018 |
| Mycolicibacterium | -0.05 | 0.05 | 0.3309 | 0.5459 |
| Bifidobacterium | -0.07 | 0.09 | 0.4182 | 0.6572 |
| Youngiibacter | 0.02 | 0.03 | 0.4541 | 0.6811 |
| Bacillus | -0.03 | 0.04 | 0.5138 | 0.7372 |
| Paenibacillus | 0.01 | 0.02 | 0.5460 | 0.7412 |
| Bacteroides | 0.03 | 0.06 | 0.5781 | 0.7412 |
| Phocaeicola | 0.03 | 0.06 | 0.5840 | 0.7412 |
| Prevotella | 0.02 | 0.05 | 0.6939 | 0.8481 |
| Saccharomonospora | 0.02 | 0.40 | 0.9696 | 0.9696 |
| Streptomyces | -0.01 | 0.04 | 0.8831 | 0.9696 |
| Ruminococcus | -0.01 | 0.03 | 0.8775 | 0.9696 |
| Aristaeella | -0.01 | 0.04 | 0.8411 | 0.9696 |
| Thermoactinomyces | 0.03 | 0.40 | 0.9351 | 0.9696 |
| Carnobacterium | -0.02 | 0.40 | 0.9518 | 0.9696 |

**Figure 19.**
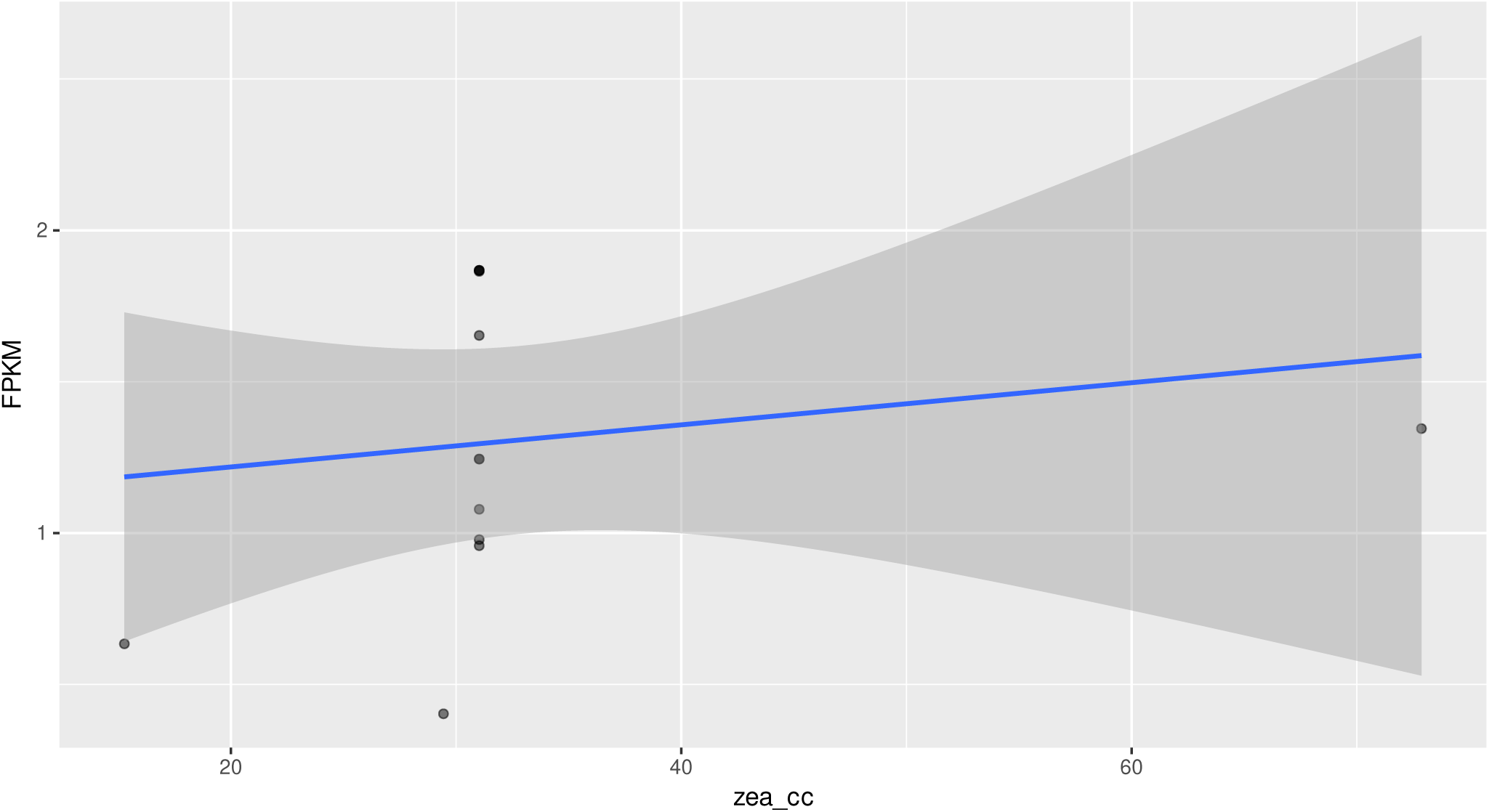
Relationship between ZEA concentration (zea_cc) and macrolide resistance gene counts (FPKM). The scatter plot shows individual data points for ARG FPKM values across a range of ZEA concentrations, with a fitted linear regression line (blue).

**Figure 20.**
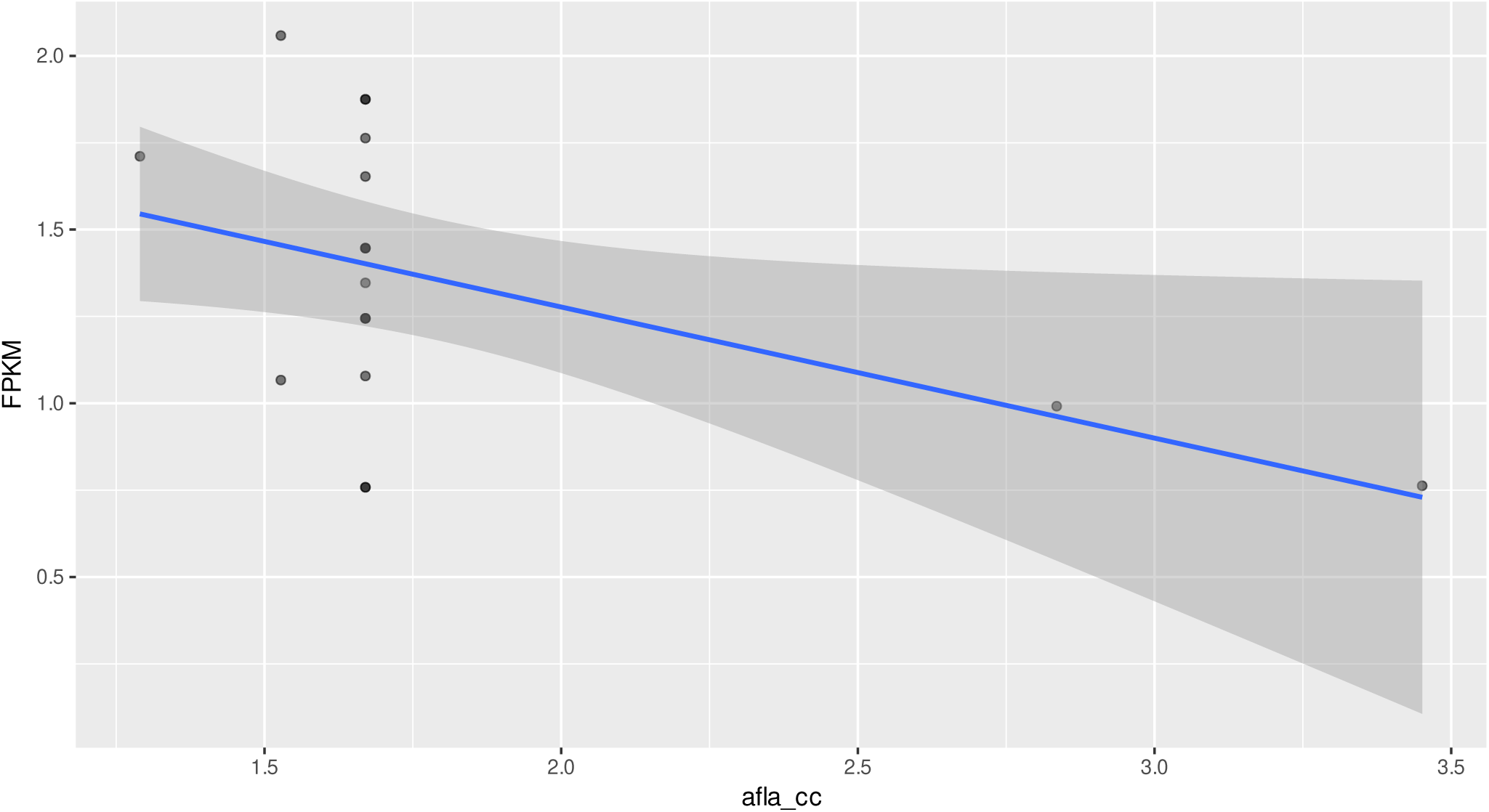
Relationship between AFs concentration (afla_cc) and ARG counts (FPKM). The scatter plot shows individual data points for tetracycline resistance gene FPKM values across a range of AFs concentrations, with a fitted linear regression line (blue).

**Table 16.** Linear model results of specific ARG FPKMs and units of the affecting conditions. ARGs affecting macrolides were tested by ZEA, and ARGs affecting tetracyclines were tested by aflatoxin.

| Model | $\beta$ | Std. Error | p-value |
| --- | --- | --- | --- |
| fpkm ∼ zea_cc | 0.006957 | 0.011491 | 0.5572 |
| fpkm ∼ afla_cc | -0.3774 | 0.1708 | 0.0411 ** |

